# Optimizing the connectivity of protein conformations to untangle ensemble refinement

**DOI:** 10.64898/2026.09.17.752399

**Authors:** Spencer K. Passmore, James M. Holton, Nadia A. Zatsepin, Andrew V. Martin

**Affiliations:** Department of Physics and Astronomy, Swinburne University of Technology, Hawthorn, Victoria 3122, Australia; Department of Biochemistry and Biophysics, University of California, San Francisco, CA 94158-2330, USA; Molecular Biophysics & Integrated Bioimaging Division, Lawrence Berkeley National Laboratory, Berkeley, CA 94720, USA; Stanford Synchrotron Radiation Lightsource, SLAC National Accelerator Laboratory, Menlo Park, CA 94025, USA; School of Science, STEM College, RMIT University, Melbourne, Victoria 3000, Australia

## Abstract

Proteins naturally adopt multiple conformations in mediating cellular processes, and ensemble models are used to fit X-ray crystallography data that captures this heterogeneity. In practice, ensemble refinement produces only minor improvements in agreement with experimental data (*R*_free_) over single-conformation models. It has recently been shown that ensemble models are universally trapped, or “tangled”; refinement algorithms strain each individual conformation in the model to fit the electron density in its immediate vicinity, missing more harmonious ways to arrange the collection of protein conformations to fit the electron density. Here, we demonstrate that this type of trap may be escaped by formulating the construction of low-energy conformations from individual conformer coordinates as an integer linear programming problem. The method successfully recovered the two original protein conformations from a previously published synthetic dataset that traps current refinement methods. Inclusion of the method in an automated refinement procedure with real data is shown to improve *R*_free_ and reduce geometric strain in a four-conformation model by comparison with controls. Applying this method in combination with human input and fitting low-occupancy waters to density features in the bulk solvent, we produce models for deposited datasets of three separate 14–19 kDa proteins with greatly improved geometry and *R*-factors. This includes a 0.77 Å six-conformation model of the SARS-CoV-2 macrodomain Mac1 (PDB ID: 44PS) with an *R*_work_ of 4.7% and an *R*_free_ of 6.4%.

## I. Introduction

The biomechanics underlying functional protein interactions with other molecules are the basis of biology, and their understanding is vital to medicine. These interactions are frequently enabled by the ability of a protein to undergo internal structural changes, which may be global or localized to specific regions [1, 2]. This property of variation in structure between otherwise identical proteins, representing multiple possible conformations for the amino acid sequence, is referred to as heterogeneity.

X-ray protein crystallography is used to capture the diffraction pattern of a protein crystal, which encodes the average structure of an ensemble of proteins. The word “average” is key here; as the molecules in the ensemble typically occupy a multitude of distinct conformations at any given point in time [3]. These may be representative of their states during physiological processes [4–6], as well as heterogeneity due to crystal-packing effects [7]. This conformational variation is further magnified by coordinated multi-atom thermal motions of proteins [8–10], which is of particular relevance to room-temperature experiments [5, 7, 11].

When working with X-ray data, models of proteins as ensembles can offer better representations of the underlying structural signal over traditional single-conformation models [1, 4, 12–14]. In an ensemble model, each atom is assigned an alternate location (altloc) for each conformation. All atoms in the conformation are given a fractional occupancy value that models the fraction of proteins in the crystal that take up said conformation. The hybrid of the single-conformation and ensemble approaches is a multi-conformer model, where the number of altlocs used varies across the model [15]. These approaches can offer crucial insights into dynamical protein interactions. For example, one study [16] considering 10,000 models deposited to the Protein Data Bank (PDB [17]) found over a third were best described using multiple distinct binding site conformations; though most such cases were missed due to being modeled in single conformations.

The accuracy of a protein model derived from X-ray crystallography is assessed by many metrics that judge the fit to the X-ray data and the physical plausibility of the structure [18]. One of the key metrics is *R*_free_, which measures the accuracy of the structure against X-ray intensities kept hidden from the refinement algorithm. Simulation work by Holton *et al*. [13] estimates the lower bound imposed by noise on *R*_free_ is as low as 5%, similar to the values achieved by small-molecule crystallography. Holton *et al*. propose two physical complications that a model must address to reach this lower bound.

First, a model must have a treatment for the non-flat bulk solvent, such as by using solvent-density distributions informed by molecular dynamics simulations [19]. Second, a model must, to some degree, represent the conformational ensemble contained in the crystal. However, in practice, ensemble and multi-conformer models produce only slight improvements in *R*_free_ [13, 20]. This result was most clearly demonstrated in a recent initiative to re-refine over 60,000 PDB models as multi-conformer models using `qFit`, which saw a median decrease in *R*_free_ of 0.01 [14].

It has recently been recognized that ensemble refinement algorithms are unable to escape “density misfit barrier traps” that arise from atom altlocs being assigned to the wrong conformation [20]. This phenomenon is also referred to as tangling, because its impact on refinement often has the appearance of ‘tying’ conformations together when they are viewed overlaid in visualization software. The Bragg reflections measured in X-ray crystallography encode the *average* electron density across all protein conformations in the sample over time and space. However, by measuring the average density, the information necessary to link together the parts of the density contributed by an individual physical conformation is lost. This “conformation blindness” creates an inherent ambiguity for ensemble refinement – which parts of the density should each conformation fit? Broadly speaking, model conformations are anchored by the *local* electron density during ensemble refinement, irrespective of whether that density corresponds to a real contiguous conformation of the original sample. If the conformations are not excessively strained, as judged by empirical geometric rules, they are unlikely to be freed from a mixed-up fit by current refinement algorithms [20]. Even under highly optimistic assumptions, the sheer number of atoms in macro-molecules affords almost countless ways for model conformations to be misfit – with tolerable strain – to segments of electron density that in reality represent atoms belonging to multiple, structurally distinct conformations of the original sample. As such, ensemble refinement converges on higher-energy model conformations that mix and warp the local geometric conformations of real conformational states in the sample.

Hopkins *et al*. [20] present a synthetic 1 Å dataset generated from a protein ensemble model with a highly idealized geometry. This represents a plausible “ground-truth” structure which a reliable refinement method would be expected to converge on if it was not trapped by tangling. The ensemble was deliberately simple, containing two equal-occupancy conformations within a flat bulk solvent. Further, these facts about the ground-truth were taken as known. Frontier refinement algorithms, including `phenix.refine` [21], `refmac` [22], and `qFit` [15, 23, 24], all converge to models with significantly worse geometries and poorer agreement with reflection data than the “best possible” model; even with simulated annealing [20]. This issue does not occur in analogous, single-conformation scenarios with synthetic data. These findings have solicited the development of new algorithms; though success has so far been limited to simpler cases where the initial model contains just one trapped atom or side chain [25].

This article places its focus on resolving three kinds of traps unique to ensemble refinement, each involving a different kind of “mix-up” in the model components attributed to the electron density. These are conformation mix-ups (density misfit barrier traps), coordinate mixups, and identity mix-ups – as summarized in Table I. Section II introduces a method to escape conformation mix-ups, wherein finding the minimum-energy conformational ensemble that can be constructed from a set of alternate locations (altlocs) is formulated as a binary integer linear programming (ILP) problem. This formulation is then extended to handle coordinate mix-ups in 2-conformation ensembles. Section III reports the successful application of the method to the synthetic scenario constructed by Hopkins *et al*., using an automated algorithm that fully untangles the trapped 2-conformation model. Section IV demonstrates the approach improving the geometry and *R*_free_ of small ensemble models of real data. Substantially improved models for previously deposited high-resolution datasets are attained by applying a procedure designed to reduce conformation mix-ups and fit weak solvent density features. The results additionally highlight the prevalence of identity mix-ups at regions of overlapping macromolecule and solvent density, indicating an outstanding challenge for application to real datasets.

**TABLE I.**
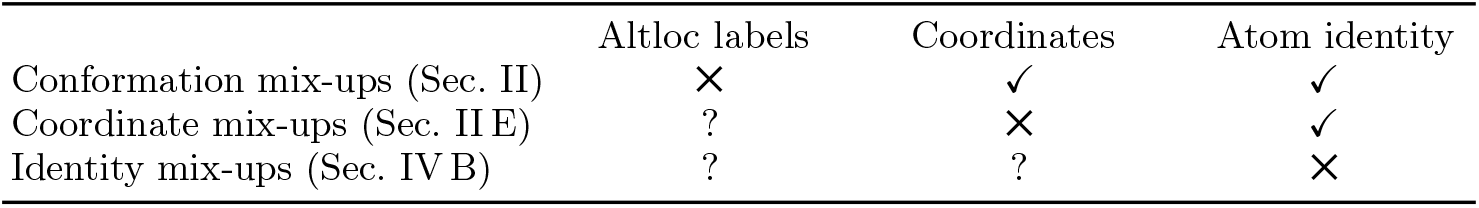
Classification of mix-up traps in ensemble refinement. Each trap directly results from an error in a distinct model component (crosses). In conformation and coordinate mix-ups, other model components are unaffected or only slightly deviated (ticks). Question marks indicate cases where a component may be wrong as a consequence of the mix-up.

## II. Method – alternate location optimization

### A. Background

Structure determination methods seek to fit the structural signal encoded in X-ray data with low-energy, geometrically plausible structures. In general terms, algorithms seek to minimize

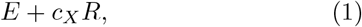

where *E* is the statistical “geometry energy”, a measure of the overall deviation of the geometric relationships from ideal library values, and *R* measures the disagreement with the X-ray data. The term *c*_*X*_ is a weight parameter.

A multitude of geometric relationships in the structure are considered by refinement programs when calculating *E*, including

- bond lengths,
- planarity,
- chirality, and
- bond angles,
- dihedrals (torsion angles),
- Van der Waals (VDW) forces.

Refining even a small protein model (*∼* 5 kDa) typically involves considering thousands of covalent geometries and tens of thousands of VDW interactions. These are compared to a library of semi-empirical ideal values and standard deviations, informed by both chemical rules and from structural models resolved to ultrahigh resolutions [21, 22]. Actual physical forces and fields are not computed, nor are quantum mechanical effects. The impact of the local structure on the ideal value for a specific type of geometry may still be accommodated with, for example, the conformation-dependent library in `Phenix` [26]. Ideal geometries are largely consistent across contemporary structure determination software packages [27].

A particular atom may occupy different coordinates depending on the conformation a protein takes. In places where the electron density clearly represents multiple conformations, atoms are traditionally assigned two or more altlocs and corresponding “occupancies”. The occupancies are fractional values which represent the proportion of asymmetric units in which the atom occupies the altloc coordinates.

For this work, it is necessary to maintain a clear distinction between an atom, conceptualized as a single entity across all conformational states, and the altlocs it may occupy. For this reason, this work refrains from using the term “conformer” – which while commonly used, refers to an atom as it exists in an individual conformation. Additionally, for synthetic data, this work defines the “correct” choice of altloc conformation labels as the one that minimizes the average RMSD of the model conformations with the synthetic ground-truth conformations. An incorrect bond is one which connects altlocs that, based on this definition, should be in different conformations.

An example of tangling is a single-atom conformation mix-up [20]. In this trap, the altloc coordinates of the atom are well-supported by the electron density encoded by the X-ray data. However, unnecessary strain in the geometry arises from how the conformations are arranged.

A single-atom conformation mix-up is illustrated for two conformations in Fig. 1. In the tangled state (Fig. 1, left), altloc A and altloc B are at coordinates that provide a close match to the electron density. However, the geometry is strained, and in tension with the X-ray term.

**FIG. 1.**
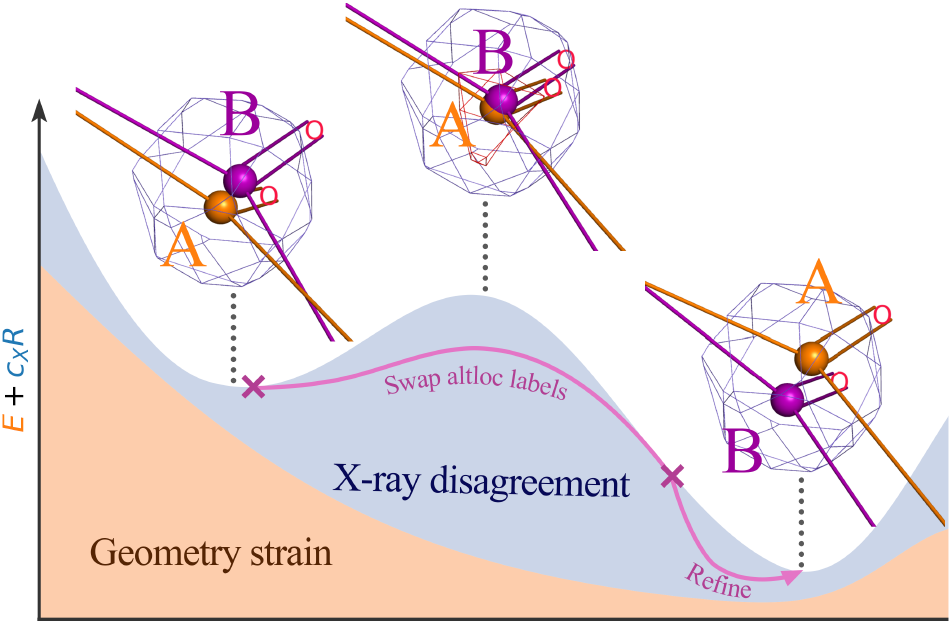
Mechanism of single-atom conformation mix-ups [20]. The insets show a modeled atom trapped in a conformation mix-up (left), the same atom when freed from the trap (right), and the intermediate of the two states (center). The lower plot illustrates the relative size of the geometry (*E*, orange) and X-ray (*c*_*X*_ *R*, blue) penalty terms that refinement software computes for each model. In the left model, the atom altlocs strain the geometry of their respective conformations, but are in good agreement with the conformation-agnostic X-ray data. This corresponds to a local minimum in the refinement cost-function *E* + *c*_*X*_ *R*. The conformations are considered to be “mixed up” because they can be rearranged to occupy similar coordinates with reduced strain (right).

It is important to appreciate that the conformations are not literally entwined; they occupy separate volumes in real space. Instead, the geometric strain is a result of each model conformation fitting electron density that the *other* conformation could fit with more ideal geometry. To fix the model with a series of small steps, as done in ordinary refinement, would require moving the coordinates away from their good fits to an intermediate state where they have strong disagreement with the X-ray data (Fig. 1, center). Swapping the altlocs untangles the model conformations, so that subsequent refinement relaxes them to aligned local minima in the geometry and X-ray terms – improving both measures (Fig. 1, right).

A conformation mix-up involves a single atom only if all covalent bonds with the atom are incorrect. Fig. 2 shows an example of a trap where this is not the case. An incorrect bond between atoms 2 and 3 means that atoms 1 and 2 of each conformation is stitched together with atoms 3, 4, and 5 of the other. While ordinary refinement is unlikely to fix a single-atom conformation mixup, here it is nearly impossible – multiple atoms would need to “pass through” each other to correct the problem. However, switching the connections between altlocs at the incorrect bond – by swapping the conformation labels of all altlocs either before or after it – escapes the trap. Notice how reconnecting the atom altlocs in this way alters fewer bond lengths and angles compared to swapping the altloc labels of atom 3. This suggests the connectivity of altlocs is, in some senses, more fundamental to conformation mix-ups than the altloc labels assigned to each individual atom.

**FIG. 2.**
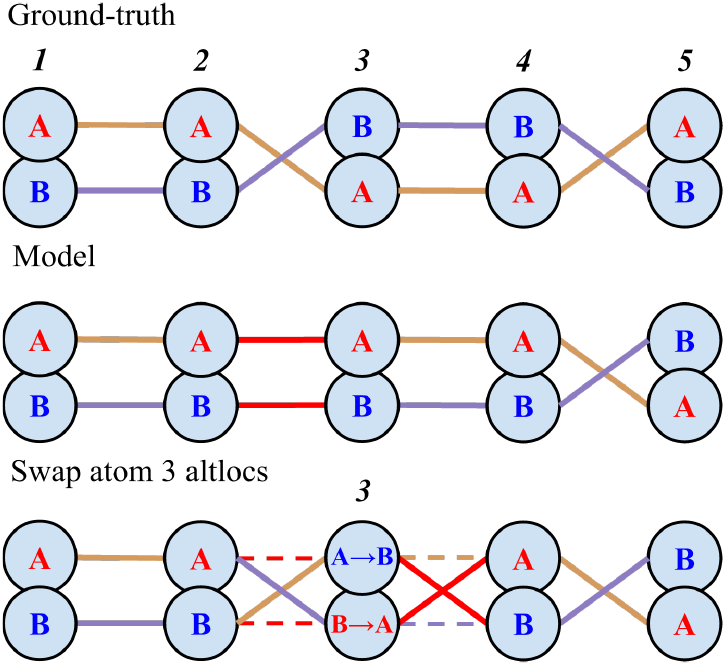
Schematic of a conformation mix-up caused by a single incorrect bond. Top: A 2-conformation structure. Middle: A model of the structure with a single incorrect bond between atoms 2–3. Bottom: Model after swapping the altloc labels of just atom 3 – this corrects the original issue, but results in an incorrect bond between atoms 3–4. Correcting the model requires swapping the altloc labels of all atoms on one side of the incorrect bond.

Where multiple incorrect bonds are present in the model due to conformation mix-ups, fixing one may produce worse geometries at distant residues. Fig. 3 shows an example where *E* is decreased only by switching the connections at two bonds, swapping the conformation labels of the atoms in between. This change is not obvious, as it increases the strain on the local geometry at one of the two bonds. However, switching the connection of only the obviously strained bond, and at the loop-forming disulfide bridge, strains the geometry of the bridge and produces overlaps in Van Der Waals radii at separate parts of the model. Thus a method to resolve these traps must be able to consider the impact of changes in connectivity on distant geometries in the model, and to fix multiple incorrect bonds simultaneously.

**FIG. 3.**
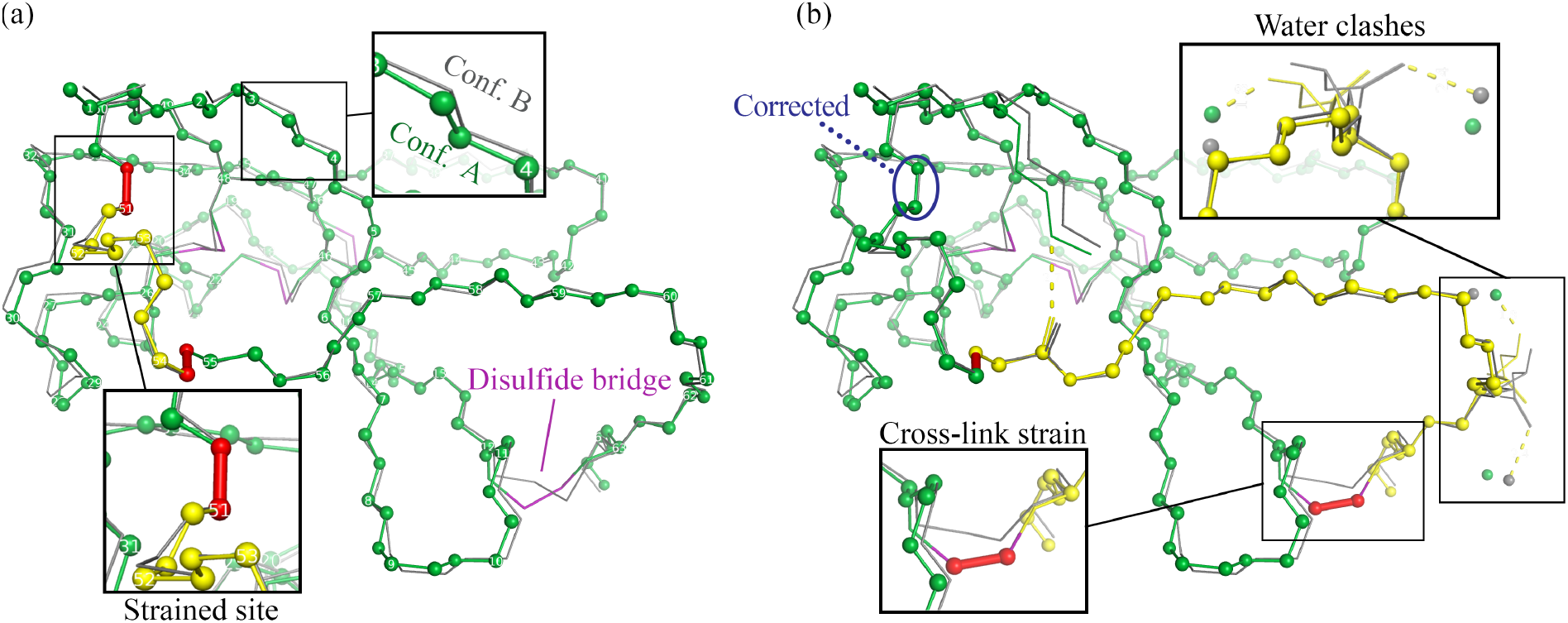
Long-range conformation-mix-up trap in a 2-conformation model caused by two incorrect bonds. Spheres correspond to atoms in the protein backbone of one conformation. Purple lines indicate disulfide bridges. Gray lines trace the backbone of the other conformation. Bonds are colored red where they connect altlocs that should be in different conformations. (a) A 2-conformation model with 4 consecutive residues assigned the wrong conformation labels. Swapping the connectivity at the two incorrect bonds would improve the overall geometry, but only from relieving strain at the depicted “strained site” of the upper incorrect bond. (b) The outcome of correcting the strained site by swapping the altlocs of all residues past its incorrect bond. The result is a model with worse geometry: close contacts (dashed lines) appear at side chains of distant residues, and the geometry of a disulfide bridge becomes strained.

### B. Constructing the problem

The geometry energy *E* [see Eq. (1)] is a weighted sum of squared *z*-scores for the geometric features of the conformations in the model, given by

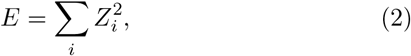

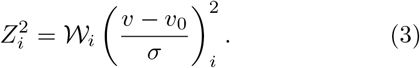

Here, *i* indexes a particular geometric measure instance (e.g. a bond length or an angle made by three atoms), *v* is the value in the model, *v*_0_ is the ideal target value, *σ* is the standard deviation, and _*i*_ is a weight parameter. In single-conformation modeling, the geometries and *E* are uniquely determined by the coordinates and their atom types. This is not the case for ensemble models. Consider a model of two conformations with equal occupancy. Swapping the labels assigned to the two altlocs of a particular atom creates a new model of two conformations with the same fit to the X-ray data, but changes the geometry energy *E*. The number of distinct ensembles of conformations that can be constructed from a set of altlocs is (*C*!)^*a−*1^, where *a* is the number of atoms in the protein and *C* is the number of protein conformations in the crystal. For example, a 3-conformation model of lysozyme has over 10^1000^ unique alternative ensembles with identical coordinates. If one of these were to have a lower *E*, it would represent a more geometrically plausible model with an identical fit to the X-ray data assuming equal occupancies. Finding such alternative ensemble models, and thus lowering *E* without affecting *R*, is a process we term *connectivity optimization*.

It is worth pausing here to develop terminology for describing the construction of a conformational ensemble from a set of altlocs. Figure 4a shows a simple case where two covalently bonded atoms each have two altlocs. One can see that there are four possible bonds or bond lengths, depending on how the altloc labels are assigned. We define these possible geometries as *geomections*. When active, a geomection represents an actual physical geometry within a conformation. In Fig. 4a, there are four bond (length) geomections for the two atoms; two of which are active. Similarly, Fig. 4b shows there are eight angle geomections for three covalently bonded atoms in two conformations; again, two are active. Each geomection couples a possible geometry between altlocs to the condition the model must satisfy for that geometry to exist: the altlocs involved must be assigned the same conformation label. As such, finding the lowest-energy set of active geomections is equivalent to finding the lowest-energy assignment of altloc labels.

**FIG. 4.**
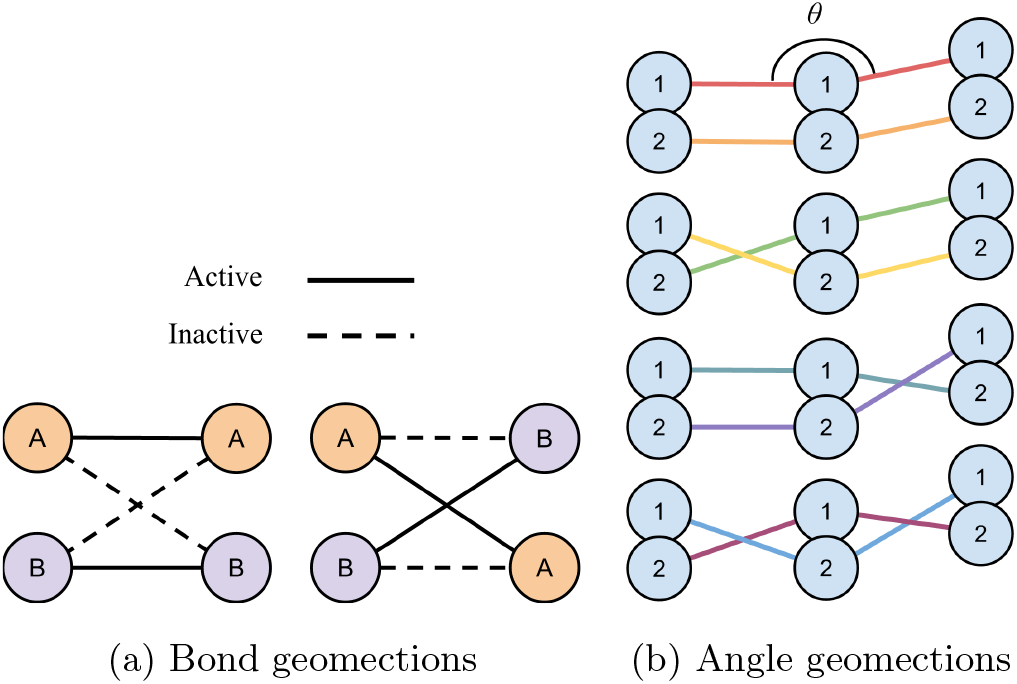
Schematic of the geomections for the covalent bond length and angle geometries in a 2-conformation model. (a) The four bond geomections for two bonded atoms in a 2-conformation model, representing the possible bond lengths across all allowed assignments of altloc labels. (b) The eight angle geomections for three covalently bonded atoms in a 2-conformation model. The number of bond geomections and angle geomections scales with the square and cube of the number of conformations, respectively.

This bears similarities to the traveling salesman problem (TSP), where the objective function to be minimized is the distance of a route that passes through a collection of cities. In the TSP, paths represent the smallest subunits of cost (distances) and are options for connecting cities together. Similarly, geomections represent the smallest subunits of cost (geometry deviations) and are options for connecting altlocs into the same conformation.

We define a binary “switch” variable corresponding to whether a particular bond geomection between atoms *m* and *n* is active,

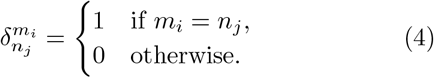

Here, the superscript and subscript terms of *δ* have the form *a*_*k*_, denoting the conformation that altloc *k* of atom *a* is assigned to. Atom *m* at altloc *i* is bonded to atom *n* at altloc *j* if and only if 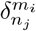= 1. Figure 5a shows the possible configurations for the bond geomections between two atoms in a model with two conformations, A and B. Atom *m* has altlocs *m*_1_ and *m*_2_, and atom *n* has altlocs *n*_1_ and *n*_2_. Non-bond geomections and clash geomections, respectively representing VDW interactions and steric clashes between pairs of atoms, are similarly each assigned a switch variable *δ*. Like bond geomections, angle geomections are active when the altlocs they involve are in the same conformation. Equivalently, it is active when the two bond geomections it ‘spans’ are both active. As such, the switch variable for an angle geomection corresponding to the angle involving altlocs *m*_*i*_, *n*_*j*_, and *p*_*k*_ is defined as a product of bond geomection switch variables 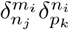. This is illustrated in Fig. 5b.

**FIG. 5.**
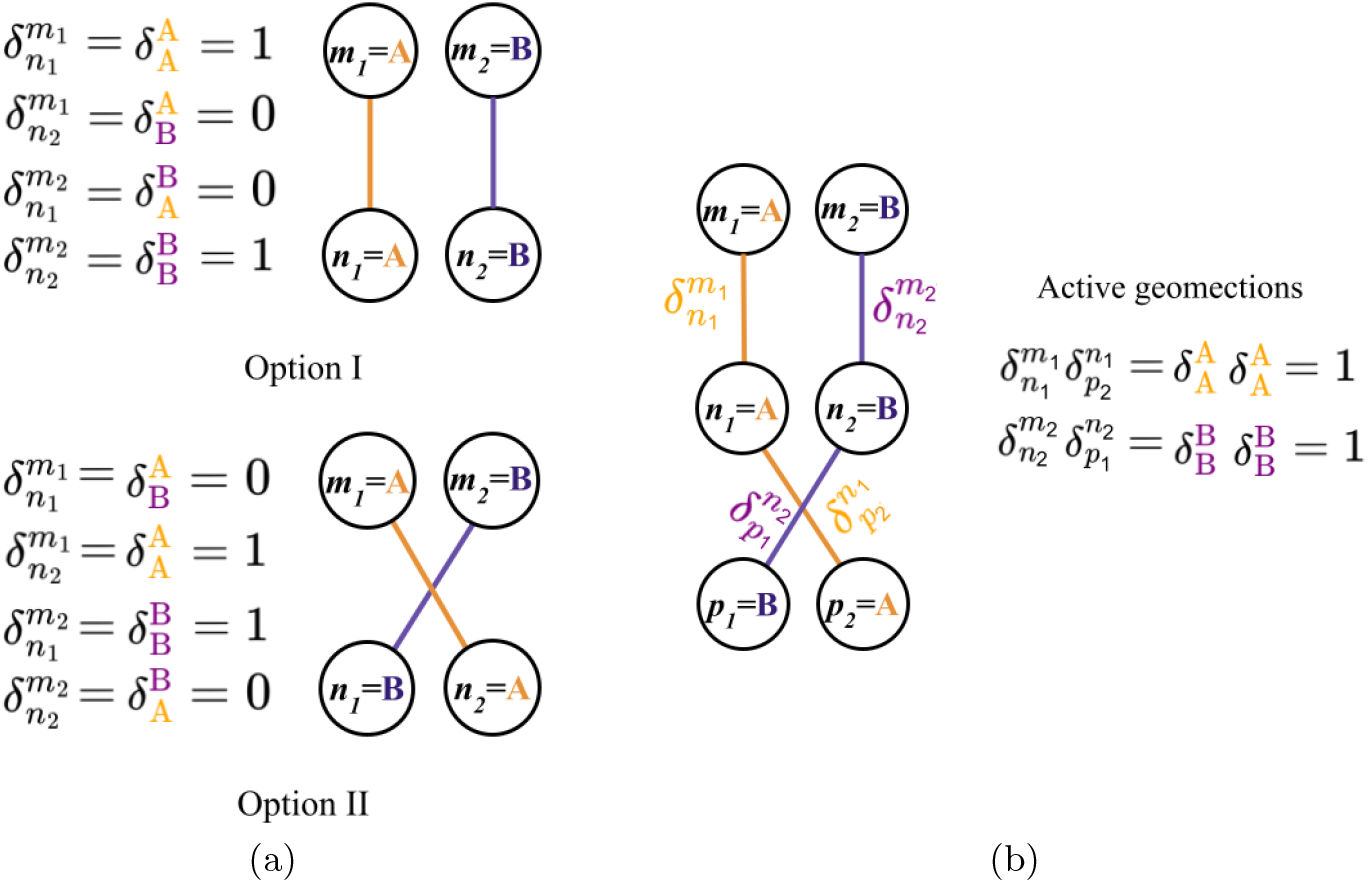
Binary switch variables for bond and angle geomections in a 2-conformation model. (a) The two physically allowed permutations for bond geomections between two atoms each with two altlocs. (b) An example of a physically allowed pair of active angle geomections between three atoms each with two altlocs. There are six other angle geomections, but these are inactive due to one or both of the involved bond geomections also being inactive. Active geomections are represented with solid lines, and correspond to (a) individual or (b) products of switch variables *c* that are “on” (equal to 1).

We may now formulate connectivity optimization as minimizing a generalized form of Eq. (2) that sums over terms representing each geomection. That is,

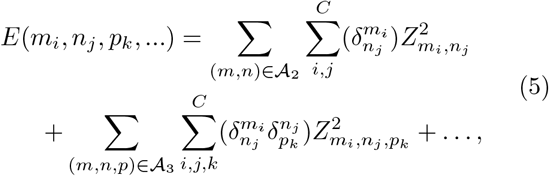

where *C* is the number of conformations and *A*_*n*_ = (*a*_1_, …, *a*_*n*_), (*b*_1_, …, *b*_*n*_), … is the collection of *n*-sized tuples of atoms corresponding to each *n*-atom geometry. This equation reduces to the simpler Eq. (2) when the binary variable values are substituted. However, its purpose is to represent *E* as a function of binary switch variables *δ* that specify the altloc label assignment when the coordinates are held fixed. This equation can be extended to include geomections for covalent geometries that involve more than three atoms, such as planarity, but this was not done in this work.

### C. Formalization as an integer linear program

Finding the physically allowed set of active geomections that minimizes *E*(*m*_*i*_, *n*_*j*_, *p*_*k*_, …) is an intractable task by brute-force methods, but relatively cheap when formulated as a binary integer linear programming (ILP) problem. ILP optimization is its own branch of mathematics, and there exist dedicated ILP solvers employing sophisticated methods to rapidly find the global minimum. These solvers follow a strategy that begins with relaxing the ILP to a continuous linear problem. “Branches” of possible solutions are then explored by fixing variables to integer values and seeking feasible integer solutions, while continuously tightening the solution space to preclude non-optimal solutions.

ILPs are formulated with integer variables and linear constraints. For the remainder of Section II, equations are written in the explicit form in which they are given to an ILP solver unless otherwise stated. Terms and products of *δ*’s, like 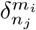 or 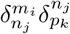, are used to represent singular binary variables in the ILP. The behavior of these binary variables is made equivalent to that described in Sec. II B using linear constraints.

Each 2-atom geomection switch 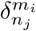is subject to the constraint that the number of geomections corresponds to the number of conformations *C*,

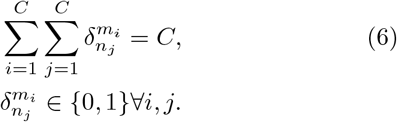

For each non-cross-link bond between atoms *m* and *n*, the constraints

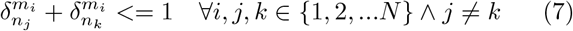

enforce mutual exclusivity for the geomections of the bond that share an altloc.

Equations (6) and (7) together enforce that 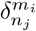 is active only when altlocs *m*_*i*_ and *n*_*j*_ are assigned to the same conformation, as in Sec. II B. Additionally, Eq. (7) ensures solutions correspond to allowed sets of active geomections, like those in Fig. 5a.

An angle geomection switch 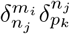 (again, representing a single binary variable) is active when the two bond geomections it spans are also active, so

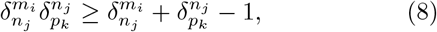

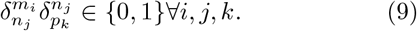

From here, there is a choice in how to proceed. For the results presented in this study we define, for each altloc, a binary variable for each conformation label that it can be assigned. However, this approach comes with a high degree of redundancy, as each geomection must track whether the atoms it is considering have been assigned the same conformation label, which requires a variable for each possible conformation the geomection could be active in. This redundancy can be eliminated by defining the problem purely in terms of the connectivity of the conformations, but this requires careful treatment for non-bond and clash interactions, and is not used here.

Proceeding with the altloc-variables formulation, each 2-atom geomection is associated with binary variables encoding the labels assigned to altlocs. A 2-atom geomection is active if its altlocs are both assigned to the same conformation, so for each conformation *X* we introduce the constraint

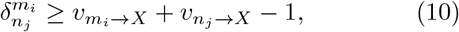

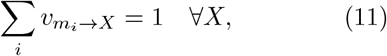

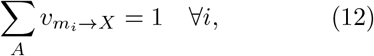

where 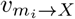 is 1 if altloc *i* of atom *m* is assigned to conformation *X*, and 0 otherwise. These constraints enforce correct label assignment behavior. A symmetry in the problem arises due to the equivalence of conformation labels. That is, we do not care whether a conformation is labeled A or B. To break this symmetry, the altlocs of the first nitrogen atom in the structure file are fixed to their original conformation labels.

### D. Implementation details

The cost function to be minimized is *E*(*m*_*i*_, *n*_*j*_, *p*_*k*_, …) [Eq. (5)]. The *W*_*i*_ values used correspond to the weights used in `phenix.refine` [21] (which may be generated by the `phenix.pdb interpretation` command). The weight on angles was scaled by a factor of 8 *×* 10^2^, as `Phenix` places a weight on bonds orders of magnitude higher than on angles.

Clashes indicate physically impossible proximities between pairs of atoms. These are instances where two atoms that do not form hydrogen bonds, and which are separated by at least three covalent bonds, have overlaps in VDW radii of at least 0.4 Å [28]. Clashes do not have a well-defined *z*-score. In its place, we impose an *ad hoc* penalty of

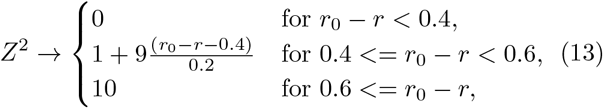

where *r* and *r*_0_ are, respectively, the distance between the two atoms and the sum of their VDW radii, in Ångstroms. This metric prioritizes resolving clashes with greater overlaps, though past 0.6 Å (which is taken as a somewhat arbitrary threshold for near certainty of a serious model error) all are penalized equally. The penalty was down-weighted for water–protein (0.5), H–protein (0.25), water–water (0.1) and H–H (0.1) clashes, under the expectation that these would be more likely to be resolved by restrained refinement than non-H protein clashes. Recall that geomections represent “possible geometries”. In contrast to other geomection kinds, it is ideal for all clash geomections to be inactive. Clash geomections provide the information necessary for the connectivity optimization to avoid introducing new clashes – particularly those that enable only minor improvements in other geometries – and to reduce the number of clashes when it is possible to do so while maintaining a reasonable geometry. Clashes were computed according to the table of VDW radii used by `molprobity` [28– 30]. These radii do not exactly match the radii used for the non-bond terms, which are taken from the CCP4 monomer library [22] (and used by `phenix.refine`). The different ideal values are, however, consistent with the weighted energy metric employed by Hopkins *et al*. [20] for model validation. A much higher weighting is applied to clashes than non-bonds; thus, where clash geomections exist between altlocs, they effectively override the CCP4 monomer library values.

Crystal-packing contacts (non-bond interactions between separate asymmetric units) have ideal non-bond separations that differ from those within the protein. For each pair of altlocs, the optimizer calculates 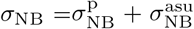, where 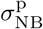 is the standard deviation from the ideal non-bond length and 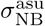 is that of the nearest crystal-packing contact. Crystal-packing contacts between unit cells were not considered when working with experimental data. In real crystals, crystal-packing contacts occur between different conformations; thus incorporating such contacts would require, in some approximate form, a model of the conformational relationships between neighboring asymmetric units. Contacts within the unit cell were still considered.

Hydrogens were treated differently from other atoms, as their weak contribution to X-ray reflections makes them particularly likely to see significant displacements from their ground-truth coordinates. First, covalent bonds with hydrogen were forbidden from changing. This means the changes to altloc labels of atoms are copied by any ‘riding’ hydrogens. Second, angle geomections involving hydrogen, while allowed to change, were not included in the cost function of the connectivity optimizer [Eq. (5)]. Testing on the synthetic dataset generated by Hopkins *et al*. [20] showed that considering angles involving hydrogen could “lock in” suboptimal geometries in the original model. See Fig. 13 in App. E for an illustrated example of this occurring in an alanine side chain.

In application to the synthetic dataset (Sec. III), a method was implemented to generate next-best solutions by forbidding solutions previously found. This feature is useful because less-optimal solutions often relax to better models after refinement.

The connectivity optimization ILP was implemented using the Python `PuLP` package for linear and mixed integer programming. By default, this package uses the open-source ILP solver `COIN-OR` [31] to compute solutions. `PuLP` supports many other well-known ILP solvers, though they require separate installations. For the results in this work, `CPLEX 21.1.1` [32] was used as the ILP solver.

### E. Coordinate mix-ups and untwist moves

In working with the synthetic dataset generated by Hopkins *et al*. [20], we identified cases where altloc coordinates were significantly displaced from the groundtruth coordinates but unaffected by refinement or connectivity optimization. Occurring significantly less frequently than conformation mix-ups, these archetypically presented with the two equal-occupancy altlocs of an atom each along the minor axis of a density ellipsoid created by two ground-truth altlocs on the major axis (Fig. 6a), in conjunction with a plausible geometry (Fig. 6b). We refer to these as *coordinate mix-ups*.

**FIG. 6.**
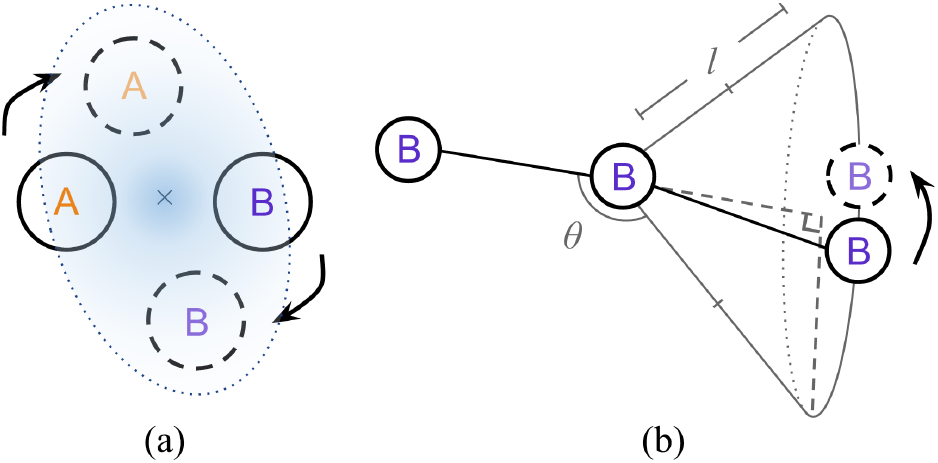
Schematic representation of untwist moves made to correct coordinate-mixups in 2-conformation models. Solid outlines correspond to model altlocs, while dashed lines correspond to the ground-truth altlocs. In this trap, the altloc coordinates are incorrect, but are caught in a local minimum with negligible tension between the X-ray data and geometric data. (a) The minimum in the X-ray data is possible only if the ground-truth coordinates approximately lie in the same plane and share the same midpoint as the trapped atom. The shaded area represents the ground-truth electron density (for illustrative purposes, the ellipticity is exaggerated). The X-ray gradient is weak or negligible near the minor axis of the ellipsoid. (b) Conformation of the trapped atom (right) and an angle it forms with two atoms (left). The trapped coordinate forms a bond length (*l*) and angle (*θ*) that is similar to those formed by the ground-truth coordinate. Where a potential coordinate mix-up is detected, one or more untwist moves that could correct the mix-up are predicted and passed to the connectivity optimizer as an optional pair of replacement coordinates.

Unlike conformation mix-ups, coordinate mix-ups are generally unaffected by connectivity optimization. Recall that conformation mix-ups are characterized by tension in the X-ray (*R*) and geometry (*E*) terms of Eq. 1, resulting from altloc coordinates close to that of the global minimum model, but assigned to the incorrect conformations. If the altloc labels are allowed to change, this tension is resolved. In contrast, coordinate mix-ups occur when *R* and *E* are both in local minima, but with “incorrect” altloc coordinates that do not correspond to the global minima. Reducing *E* only by reassigning altloc labels is thus unlikely to resolve this trap.

For the purposes of the application to the synthetic dataset in this study (Sec. III), an algorithm was developed to detect possible coordinate-mixups in 2-conformation models, and predict corresponding ground-truth altlocs. These changes to the coordinates, or untwist moves, can then be passed to the connectivity optimizer with corresponding alternative geomections for the new coordinates. Fig. 6 shows what such a move looks like. See App. B for details on the procedure used to identify untwist moves. Note that this method does not handle traps involving more than 2 conformations.

Untwist move options are represented in the connectivity optimizer with binary variables. We define the binary variable 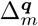 to be 1 if the altlocs of atom *m* are assigned the pair of coordinates ***q*** – representing either no change, or one of the possible untwist moves. Only one set of coordinates is allowed for each atom, which is enforced by the constraint

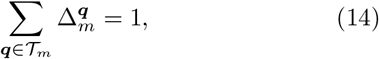

where *T*_*m*_ is the set of all allowed coordinate pairs for atom *m*, including the original coordinates.

When untwist moves are available, 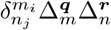 terms become the binary geomection switch variables of the ILP, replacing 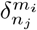 . Eq. 6 is modified accordingly, so as to remain a sum over all geomection switch variables for the geometry. Additionally, Eq. 7 is replaced with

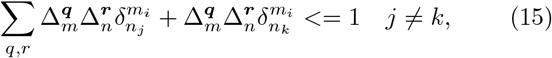

and Eq. 10 becomes

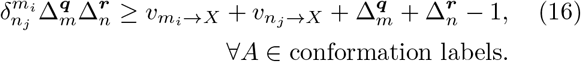

This modification transforms the 2-atom geomection cost terms in Eq. (5) as

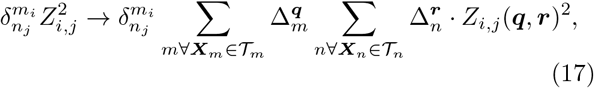

and the 3-atom terms are similarly transformed. Note that Eq. 17 represents the *δ* and Δ as separate terms, which does not follow the explicit form given to the ILP solver.

Traps affecting the connectivity of residues are more severe, as they can lead to long-range traps over multiple residues. As such, all protein backbone atoms were considered for untwist moves. Cysteine C*β* atoms were also considered for this reason, due to the cross-links they form between residues through disulfide bridges. Additionally, it was expected that coordinate mix-ups would be more reliably diagnosed in the protein backbone due to its rigidity. When working with synthetic data (Sec. III), coordinate mix-ups in side-chain atoms were observed to occur rarely relative to occurrences in the backbone atoms.

## III. Application to synthetic 2-conformation dataset

For the purpose of testing proposed methods to escape conformation mix-ups, Hopkins *et al*. present a synthetic X-ray dataset generated from a 2-conformation, equal-occupancy model of a scorpion toxin protein (PDB ID: 1aho [33]). This representative “ground-truth” model was obtained through a procedure that placed a heavy emphasis on geometry idealization. As such, the global-minimum fit to the X-ray data and geometry was expected to closely align with the model obtained by refining the ground-truth against its diffraction data. Contemporary refinement methods are capable of reaching the global-minimum fit in similar constructed scenarios where the synthetic ground-truth is a single conformation. However, in this scenario, such methods inevitably become caught in a multitude of conformation mix-ups. In idealizing the geometry of the ground-truth, the Holton weighted energy wE [20] was minimized, replacing the more conventional *Z*^2^ score as the geometry energy metric. As such, in this section, the cost function of the connectivity optimizer [Eq. (5)] was modified to have a more similar form to wE (see App. A), minus terms for geometries not considered here (dihedrals, chirality, planarity) or the worst outliers of a geometry.

The best-scoring protein model obtained by Hopkins *et al*. was used as the initial model. This “long-range traps” model was produced through rectified simulated annealing [20]; a modified method of simulated annealing in which only changes that improve the model geometry are permitted. The procedure applied to the initial model first attempts to bring the altlocs closer to those of the ground truth, regardless of conformation, by refining the model to the X-ray data without geometry restraints. This is followed by connectivity optimization to generate candidate altloc label assignment solutions. The candidate solutions are refined in parallel with geometric restraints enabled, and the best-scoring model is taken. The specific steps are as follows:

1. Refine the protein altlocs to the X-ray data without geometric restraints (unrestrained refinement) with 1 `phenix.refine` [21] macro-cycle (reciprocal-space refinement of individual altloc *B*-factors and coordinates). Solvent coordinates are unmodified.
2. Detect possible coordinate mix-ups for main chain atoms (N, C, C*α*, O) and C*β* atoms in cysteine.
3. Perform connectivity optimization with optional untwist moves (Sec. II) to generate the 25 lowest-energy altloc assignments for the entire ensemble model.
4. For each solution perform restrained refinement of protein and solvent altlocs over 6 `phenix.refine` macro-cycles. The lowest wE model is taken.

To lower computation times, connectivity optimization was constrained to exclude solutions replacing bond lengths or angles with outliers of more than 8 or 3 standard deviations, respectively. This rule was exempted for cases where the geometry contained a larger outlier in either conformation. Additionally, solution finding was performed so as to guarantee optimality within the constraints of the ILP problem, by mathematical proof. The solutions identified thus represent the lowest-energy models that can be constructed from the altloc coordinates within the constrained solution space.

An additional loop of these steps is then applied to the model resulting from the first loop, but with the search space of the connectivity optimization narrowed to focus on resolving incorrect long-distance relationships in the protein backbone and cross-links (disulfide bridges). Under the imposed restrictions,

1. C-O bond changes are forbidden unless the N-C and/or C*α*-C bond also changes.
2. Bond changes involving side-chain atoms are forbidden, excluding disulfide bridges (cysteine).
3. Multiple bond changes with backbone atoms or C*β* within a residue and its neighbours are forbidden. Atoms altered by untwist moves were excluded from this rule.
4. Untwist moves are disabled.

Individual hydrogen refinement is enabled during restrained refinement, as this was found to reduce hydrogen clashes. However, refining with individual hydrogens produces better scoring models, and was not used in the models presented by Hopkins *et al*. [20]. So that comparison with the aforementioned work remains meaningful, a single additional restrained macro-cycle is performed with riding hydrogens for all models before the lowest wE model is taken in the last step of each loop.

Figure 7 shows how the model connectivity and metrics are changed by the two connectivity optimization and refinement loops. The initial model [Fig. 7(a)] contains a large number of incorrect bonds (red), which section off segments of incorrectly labeled atoms (yellow). After restrained refinement of the best 25 solutions found by the connectivity optimizer in loop 1, the lowest wE model (Fig. 7b) is fully untangled. This model was refined from the 9th-lowest-*E* solution found by the connectivity optimizer. The model resulting from the first loop was found to be fully untangled when compared with the ground truth, with X-ray and geometry metrics (wE=18.45, *R*_free_=3.17%) close to the optimal values from “`best.pdb` ”, the model obtained from refining the ground-truth model to the X-ray data (wE=18.27, *R*_free_=3.14%) [20].

**FIG. 7.**
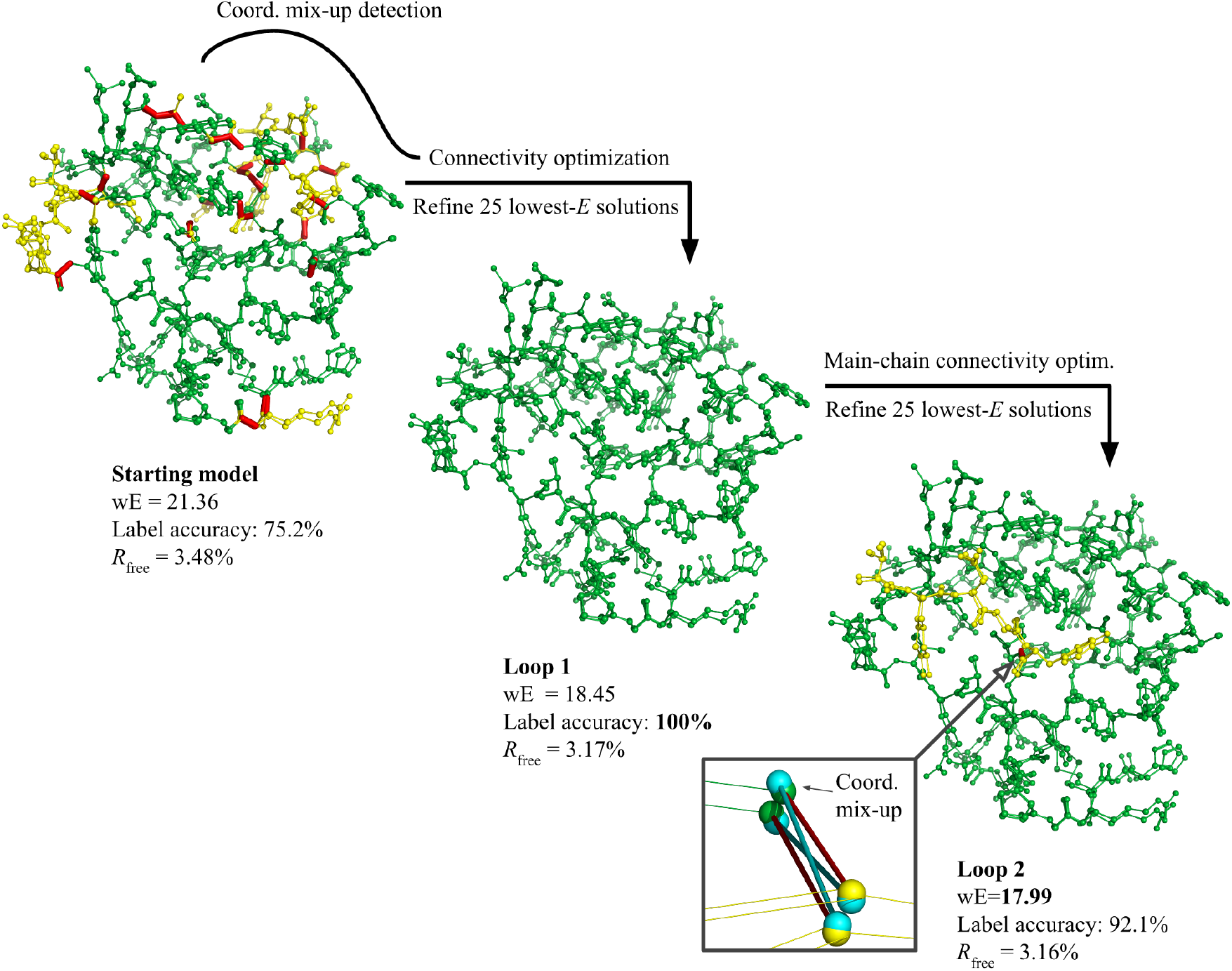
Evolution of the 2-conformation model for the synthetic scenario [20] under connectivity optimization and refinement. The two conformations of the model are shown overlaid, but they are separate in physical space. Atoms more than 0.05 Å from the ground-truth coordinates of their conformation are colored yellow. Hydrogens are not shown. Incorrect bonds (red) connect altlocs that should be in separate conformations. The label accuracy gives the percentage of atoms assigned the correct conformation labels. The bottom-right inset shows the loop 2 model at the site of its singular incorrect bond (C–N), overlaid with the ground-truth altlocs and bonds in cyan.

Fig. 7c shows how applying the modified method of loop 2 to the loop-1 model produced an “alternative hypothesis” model. The model disagrees with the ground truth, but its metrics (wE=17.99, *R*_free_=3.16%) are similar or better to those of `best.pdb` . In this model, residues 1–5 are assigned to the wrong conformations. The apparent cause is a coordinate mix-up in the N atom of the residue 6 backbone, coinciding with the incorrect residue 5–6 peptide bond (magnified in the figure) responsible for the error in connectivity. This model was the result of refining the 2nd-lowest-*E* solution found in the connectivity optimization step of loop 2. Using the connectivity optimizer, we identified an additional alternative hypothesis model where a coordinate mix-up in the Lys51 C*β* results in its side-chain conformations being incorrectly swapped. When applying the untwist detection algorithm to these models, in both cases the correction for the coordinate mix-up is among the untwist moves selected.

Resolving coordinate mix-ups was crucial to finding the untangled solution in a single loop. The untwist move detection algorithm found 143 possible replacement altloc coordinate pairs for 89 of the atoms, of which 13 were identified as consistent with the X-ray data and passed to the solver. The connectivity optimizer solution that refined to the best model included coordinate pair changes for three backbone N atoms and one cysteine C*β* atom, shown in Fig. 8. The replaced coordinates are remarkably closer to the ground truth in each case.

**FIG. 8.**
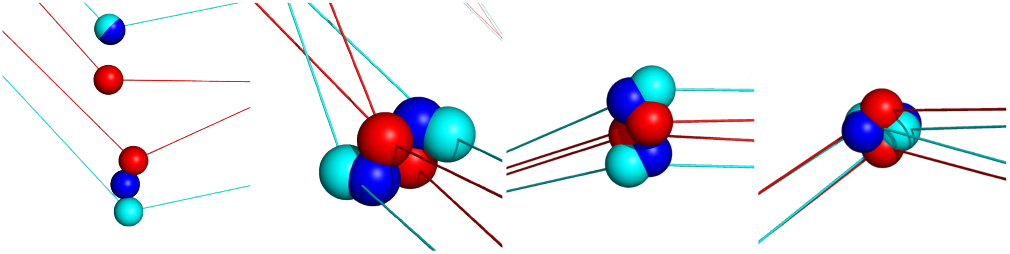
Untwist moves made in the first loop. Shown are the new coordinates selected by the connectivity optimizer (dark blue), automatically detected in the model after unrestrained refinement (red). The ground-truth model, which is unknown to the algorithm, is shown in cyan. Residues/atom names from left to right: Lys2/N, Arg18/N, Cys22/C*β*, Ser33/N. Hydrogens are not shown. Lys2/N is not truly trapped, but the algorithm allows alternative coordinates identified during untwist move detection when they have a significant separation (over 0.2 Å) that is more than twice the separation of the original coordinates. Each move brings the altlocs significantly closer to the ground-truth coordinates.

## IV. Real data

### A. Controlled test with experimental data

To test the performance of the connectivity optimizer in practice, we considered a 0.85 Å experimental dataset of dihydrofolate reductase (DHFR) [34] deposited alongside a `qFit` multi-conformer model (PDB ID: 4PSS [35], *R*_free_=16.8%) and a 250-conformation model (4PTH [36], *R*_free_=14.4%). The `qFit` model, which has 1–4 altlocs for each atom, was modified to have four contiguous conformations through copying altlocs. The anisotropic temperature (*B*) factors of the model altlocs were changed to be isotropic, as anisotropic *B*-factors may mask conformation mix-up errors [20]. The resulting model coordinates were given a random displacement averaging 0.5 Å and refined for 10 macro-cycles in `phenix.refine`.

A similar procedure was undertaken as in the first loop performed in Sec. III, with the following changes to the connectivity optimizer step:

**(I):** No untwist moves are made,

**(III):** The cost function is a sum of *Z*^2^ values [Eq. 5],

**(IV):** Instead of a batch of candidate solutions, only one solution is produced for refinement,

**(V):** solution finding was broken into a series of subproblems, so as to iterate on good but not necessarily optimal solutions within an incrementally expanding solution space (see App. C),

**(VI):** solutions were forbidden from introducing clashes between pairs of atoms that previously had no clashes, and

**(VII):** non-bond geomections were not considered.

Additionally, 10 macro-cycles were performed instead of 6 for the restrained refinement step after connectivity optimization.

The evolution of key validation measures for the model under the procedure is summarized in Fig. 9. The procedure was repeated 5 times, with starts differentiated by randomly displacing the initial model coordinates by 0.1 Å on average, then refining for 6 macro-cycles (this is separate to the random displacement and refinement of the initial model). Results for control refinement procedures without the connectivity optimization step, and with only the restrained refinement step, are also shown. Since unrestrained refinement minimizes *R*_work_ directly, one might expect that the connectivity optimization step would lead to *R*_work_ decreasing more rapidly, but no such effect is evident. However, the inclusion of connectivity optimization produces a clear improvement in *R*_free_ and geometry over the controls, up until evident overfitting past loop 13.

**FIG. 9.**
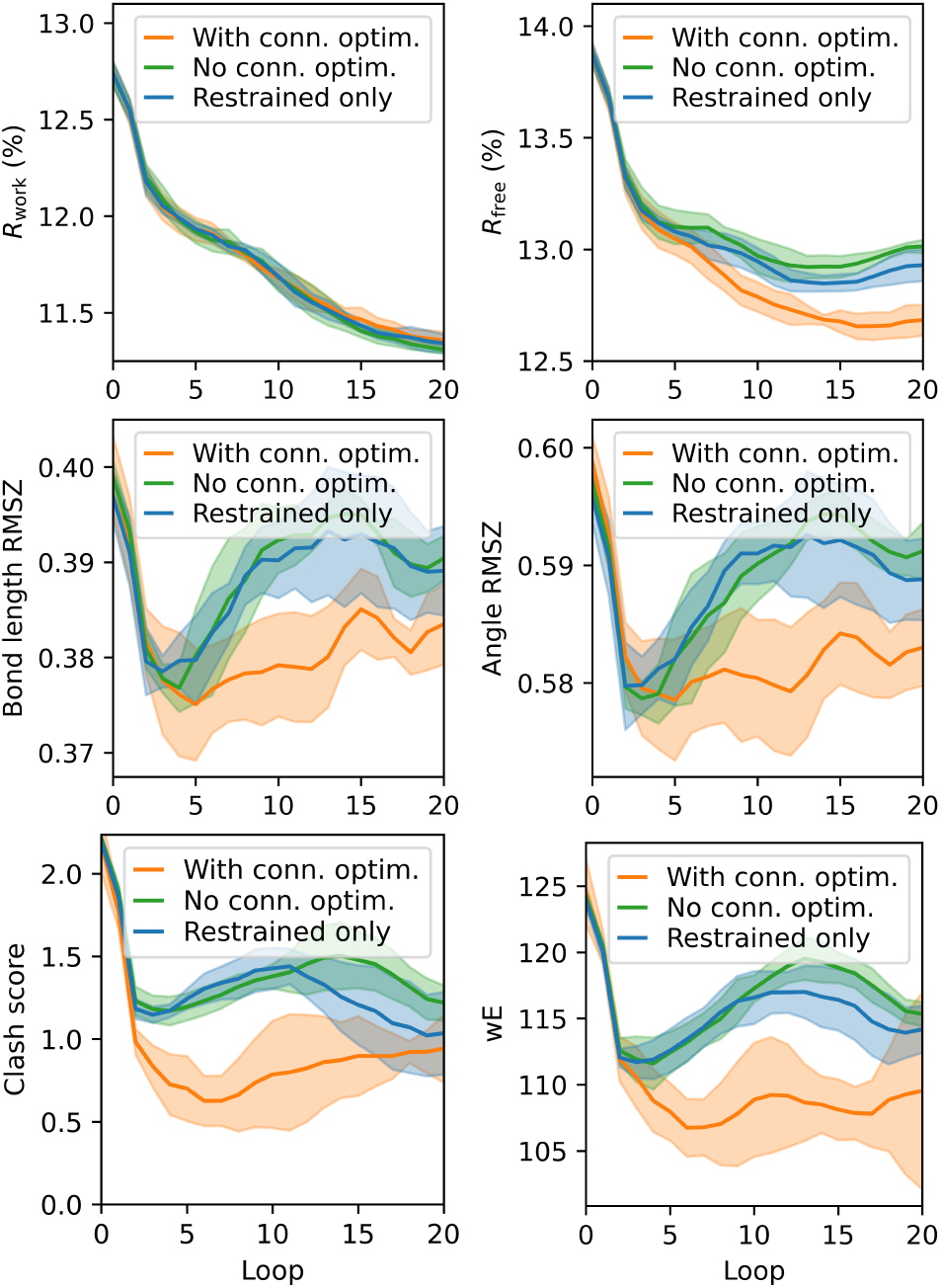
Evolution of measures for 4-conformation models of DHFR refined against experimental data [35] under the automated connectivity optimization procedure (orange) and control procedures (green and blue). The mean was taken over 5 runs, shaded regions correspond to values within 1 standard deviation of repeated runs. Each trace corresponds to a different refinement strategy for each loop: Orange – one unrestrained refinement macro-cycle, connectivity optimization, ten restrained macro-cycles. Green – one unrestrained maco-cycle, ten restrained macro-cycles. Blue – ten restrained macro-cycles. Run metrics were measured at the end of each loop on a 3-point moving average.

The clashscore values in Fig. 9 were calculated for the actual model without further changes. By default, MolProbity calculates the clashscore for the model after calling `Reduce` [29, 37, 38]. The primary purpose of this is to add missing hydrogens in optimized geometries, though here the hydrogens are already modeled. However, `Reduce` was found to make improper changes when applied to ensemble models, replacing all hydrogens and making several side-chain flips. These changes did not occur when calculating the clashscore for the same conformations individually, in separate single-conformation structure files. This behavior persisted after removing all waters, so is likely a compatibility issue with the ensemble format. Excluding `Reduce` had negligible impact on the clashscores of individual-conformation model files.

### B. Application to experimental datasets

In practice, conformation and coordinate mix-ups are not the only kind of modeling issue that can limit *R*factors. Detailed simulations of X-ray data collection suggest that, in the absence of other modeling issues, a flat bulk-solvent approximation introduces an error significant enough to limit the lowest achievable *R*_free_ value to just under 10% for 2Å models [13]. Indeed, we were unable to achieve a model with an *R*_free_ below 10% without the additional treatment for the bulk solvent described in this section. Here, we describe the process used to arrive at low-*R*_free_ ensemble models of the structure factors for high-resolution depositions – lysozyme (1.2 Å, *R*_free_=18.0% [11, 39]), DHFR (0.85 Å, *R*_free_=16.9% [34, 35]), and the SARS-CoV-2 non-structural protein 3 (NSP3) macrodomain Mac1 (0.77 Å, *R*_free_=11.7% [40, 41]). Each of these depositions are multi-conformer models produced by `qFit`.

Figure 10 summarizes the general process followed for each dataset. Six-conformation models were constructed by adding copies of altlocs to the original models, deviating the coordinates an average of 0.5 Å, then performing 20 or more restrained refinement macro-cycles. For each dataset, the initial model was first refined through an automated “connectivity refinement loop” consisting of one unrestrained refinement macro-cycle, connectivity optimization, and then ten or more restrained refinement macro-cycles. This was repeated until convergence in *R*_free_. Further repeats were then performed, with the procedure modified to add low-occupancy ordered waters, or remove them, using the conformation-aware capability introduced in `Phenix` version 2.0. All waters are assigned altloc labels corresponding to any of the six conformations in the model. After convergence in *R*_free_ was again achieved, manual adjustment of the model was subsequently performed in `coot` [42]. Changes were focused on correcting issues that could not be remediated by connectivity optimization. For example, adjusting side-chain conformations significantly displaced from density that might support them, or merging low-occupancy waters in the bulk solvent to make room for placing new waters at adjacent density features. Several repeats of this were performed, with the connectivity refinement loop, in addition to water picking and water occupancy refinement, run several times between manual adjustments. The *B*-factors of the NSP3 model were converted to anisotropic values near the end of the process. This was not done for the lysozyme or DHFR models due to substantially widening the gap between *R*_work_ and *R*_free_. See App. D for more specific details and parameters.

**FIG. 10.**
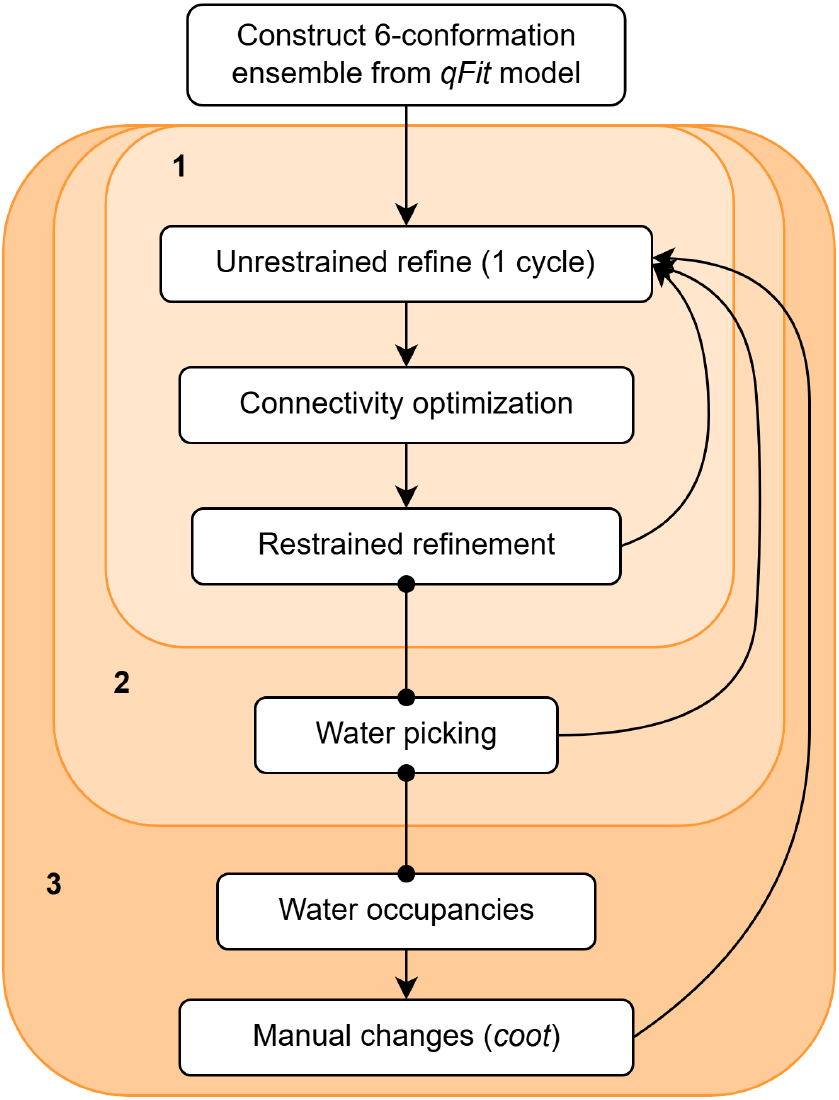
Flowchart of the procedure followed in refining the 6-conformation models (Table II). Stage 1 of the procedure (top) is the automated connectivity refinement loop used in the controlled test (Sec. IV A). Stage 2 modifies restrained refinement to perform conformation-aware water picking. Stage 3 introduces manual adjustments at the end of the loop, to fix real-space traps such as identity mix-ups. Water occupancies are also refined in this stage. Each initial model is derived from a deposited qFit multi-conformer model.

Overall, each model was subjected to hundreds of restrained refinement macro-cycles. This may be atypical for single-conformation modeling, but it appeared to be necessary for convergence here with ensemble models. The statistics of the final models are summarized in Table II. Each of these models see substantially reduced *R*-factors compared to their original depositions.

**TABLE II.**
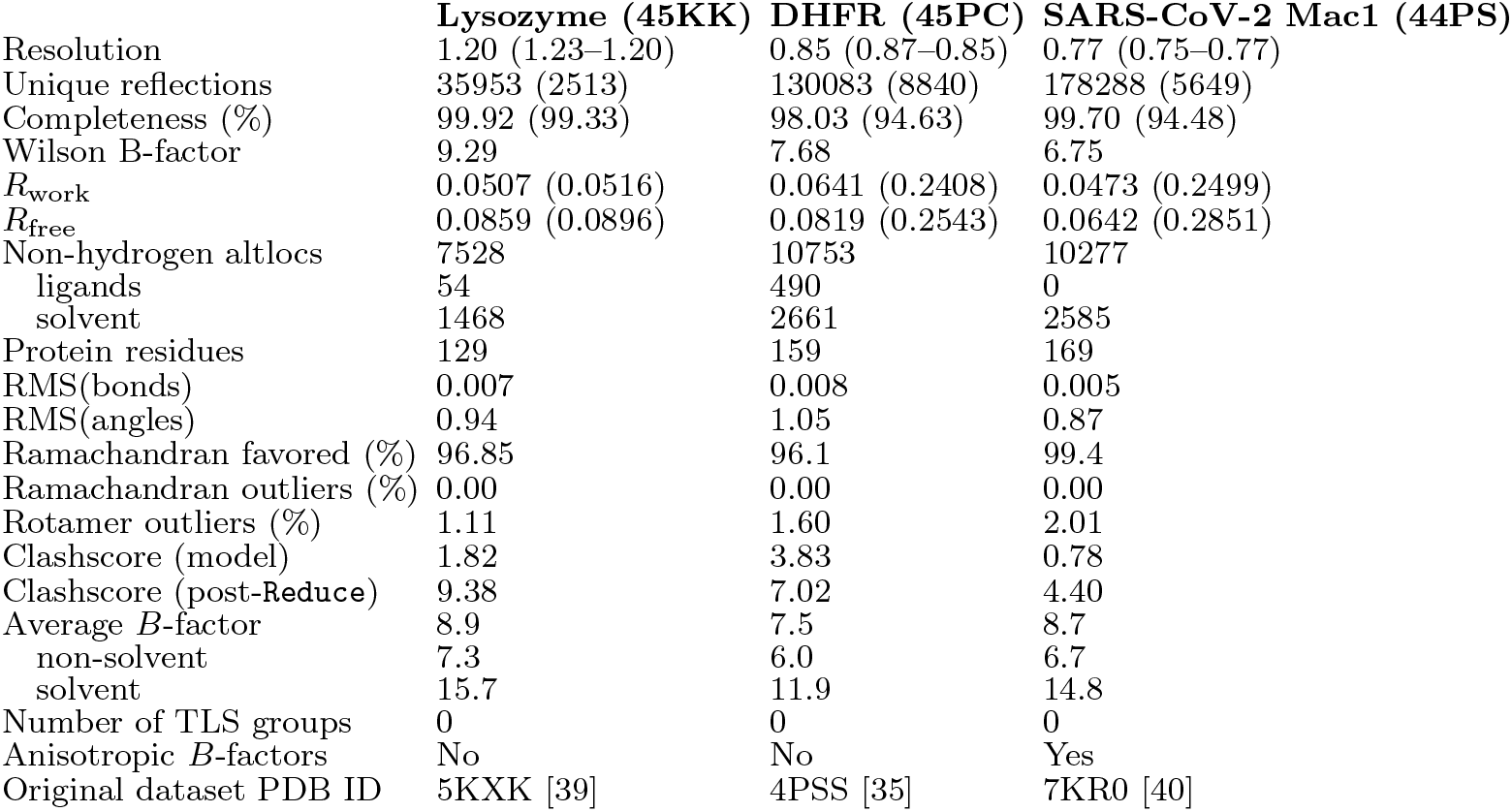
Statistics for re-refined 6-conformation models.

Before manual adjustments, identity mix-ups in the fitting of waters and protein to density features were pervasive in solvent-facing side chains. The motions of solvent-facing side chains can sweep across distances much larger than atomic separations. It is plausible – even expected – for the motion of a side-chain atom to occur in conjunction with a water or even another side-chain atom moving in to take its place; such as the case depicted in Fig. 11a. Fig. 11a illustrates how modeling the side chain and water as occupying small separate volumes across all conformations (and thus modeling an overly constrained range of motion) can provide a close fit to data that in reality comes from overlapping protein-solvent density contributions. Since the atom identities of conformation C are mixed up, the Trapped model cannot be corrected by connectivity optimization. Metrics indicating close non-bond contacts are frequently used to detect such issues in single-conformation modeling [18]. However, the reliability of this approach is diminished here, as the spread in the protein conformations allows for waters to intrude into what would be the Van Der Waals volume of a single-conformation model. Each model was improved at a number of side-chains by making manual changes to resolve apparent mix-ups. These changes were made due to being strongly supported by validation metrics – improving local geometry and the local density fit, lowering *B*-factors, and reducing *R*_free_. However, several side chains with relatively poor metrics for some conformation, and which were considered as plausible cases of complex identity mix-ups, remain untreated in each model.

**FIG. 11.**
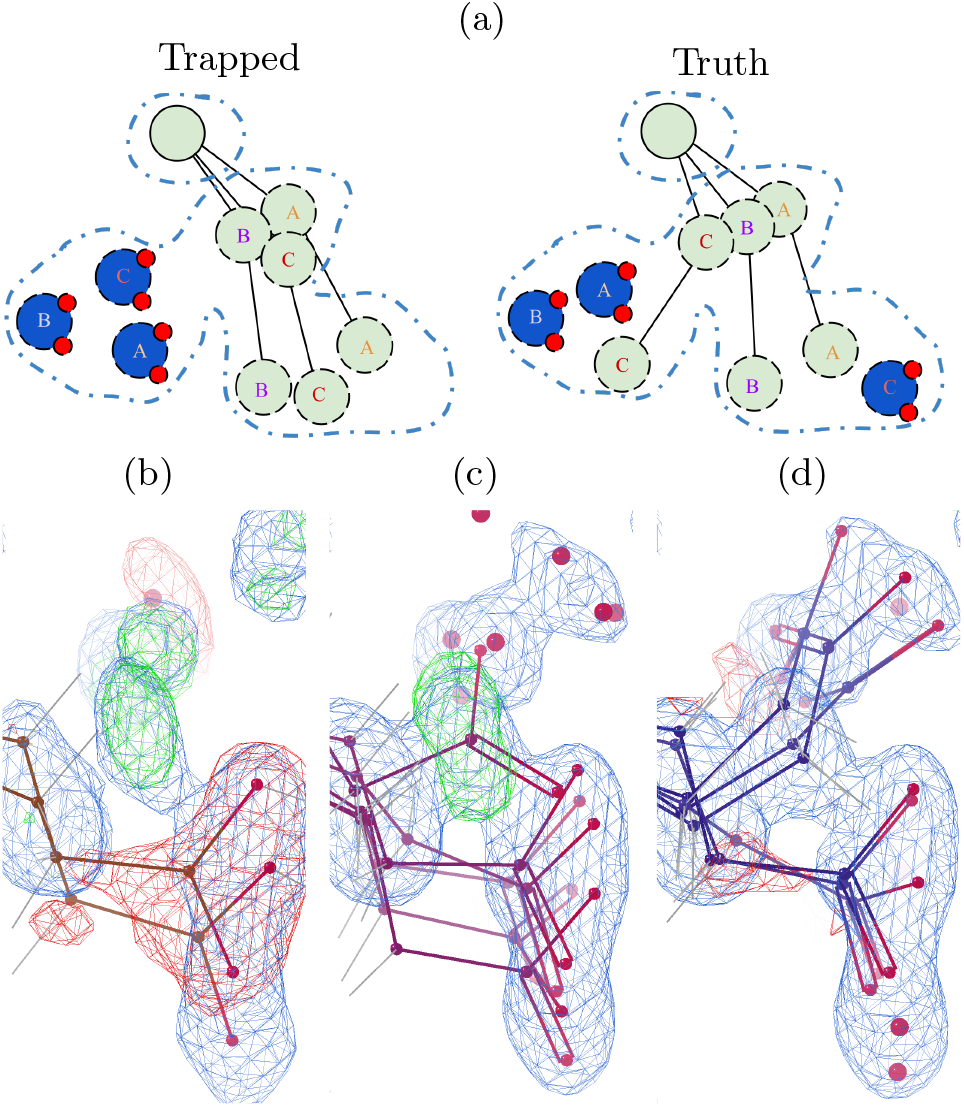
Atom identity mix-ups in ensemble models. (a) Illustrates a scenario where a model (left) incorrectly fits the average electron density of the ground-truth (right). In the Trapped model, the protein and water contributions to the density are confined to separate volumes of space. In the Truth model, the lower-left and lower-right regions of density see contributions from the protein in some conformations of the asymmetric unit, and from ordered water in others. (b– d) Show an example of the trap occurring during refinement of the DHFR model, in the folic acid ligand. (b) Original multi-conformer model (4PSS [35]). (c) 6-conformation model before manual changes. (d) 6-conformation model after manually removing waters in the upper density and rotating in conformations to replace them, and further connectivity refinement loops. FoFc maps are contoured at 3*σ*.

Figure 11b–d together show an example of a protein-solvent identity mix-up that was resolved at the folic acid ligand of the DHFR model. In the original `qFit` model (Fig. 11b), an ordered water is fitting the upper density. In Fig. 11c, the density is fit with 5 waters across 4 conformations, which have separations and angles close to the ideal geometry of the carboxyl group (R-COOH). Note how the 2FoFc map now suggests the need for an alternate rotamer to be modeled into the upper density. Additionally, the upper oxygen atoms of the upper carboxyl group conformation have abnormally high *B*-factors relative to the other conformations. After manually deleting the waters, moving the ligand conformations to fit this density in `coot`, and then refining, the resulting distinct rotamer conformation provides a better explanation for the density feature originally modeled as water (Fig. 11d). Additionally, the bottom portion of the lower-right density “blob” is now fit with waters hydrogen-bonded to the conformations of the *new* rotamer. These waters make overlapping contributions to the density with the carboxyl groups of the *original* rotamer. The conformations of the original rotamer are also more similar as a result of this change. Remarkably, the density clearly favors this interpretation of the lower region in spite of the relatively small differences between the water and ligand oxygen coordinates. A similar identity mix-up trap encompassing two adjacent side chains was also observed in this model (Fig. 14).

The models contain a large number of low-occupancy ordered waters that have the appearance of fitting high-solvent-density regions of the bulk solvent [13, 19]. Fig. 12 shows a region of the bulk solvent in the DHFR model, with the 2FoFc map contoured at a low RMSD of 0.3*σ*. This reveals the many water altlocs in this region to be well-supported by weak, elongated features in the electron density. These may serve as corrections to the flat bulk solvent model used by `phenix.refine` [43], rather than representing rare, static ordered waters.

**FIG. 12.**
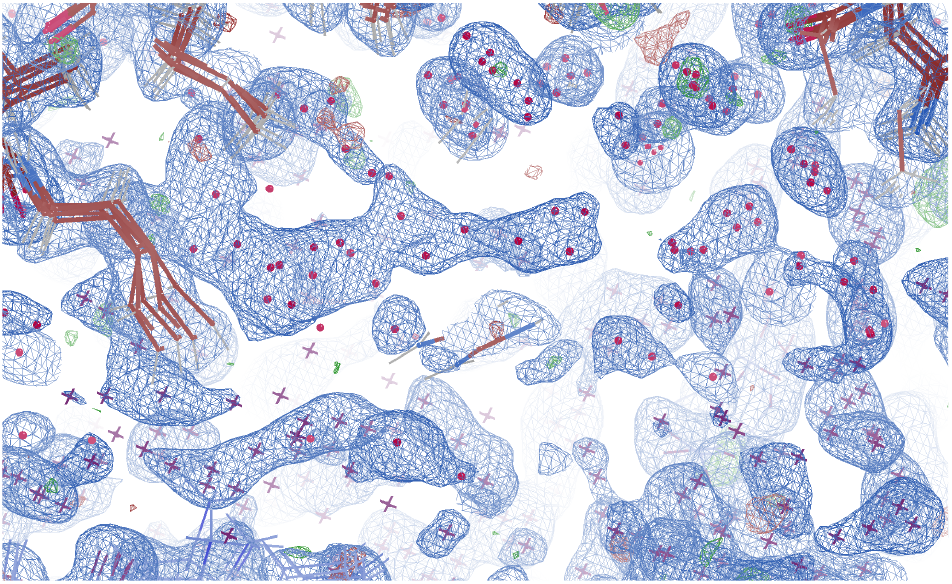
Low-occupancy waters in the interstitial solvent channels of the 6-conformation DHFR model. Dots correspond to water positions in the unit cell, crosses correspond to water positions given by the crystal symmetry. 2FoFc and FoFc maps contoured at RMSDs of 0.3*σ* (0.20 e*·*Å^*−*3^) and 3*σ* (0.29 e*·*Å^*−*3^) respectively.

**FIG. 13.**
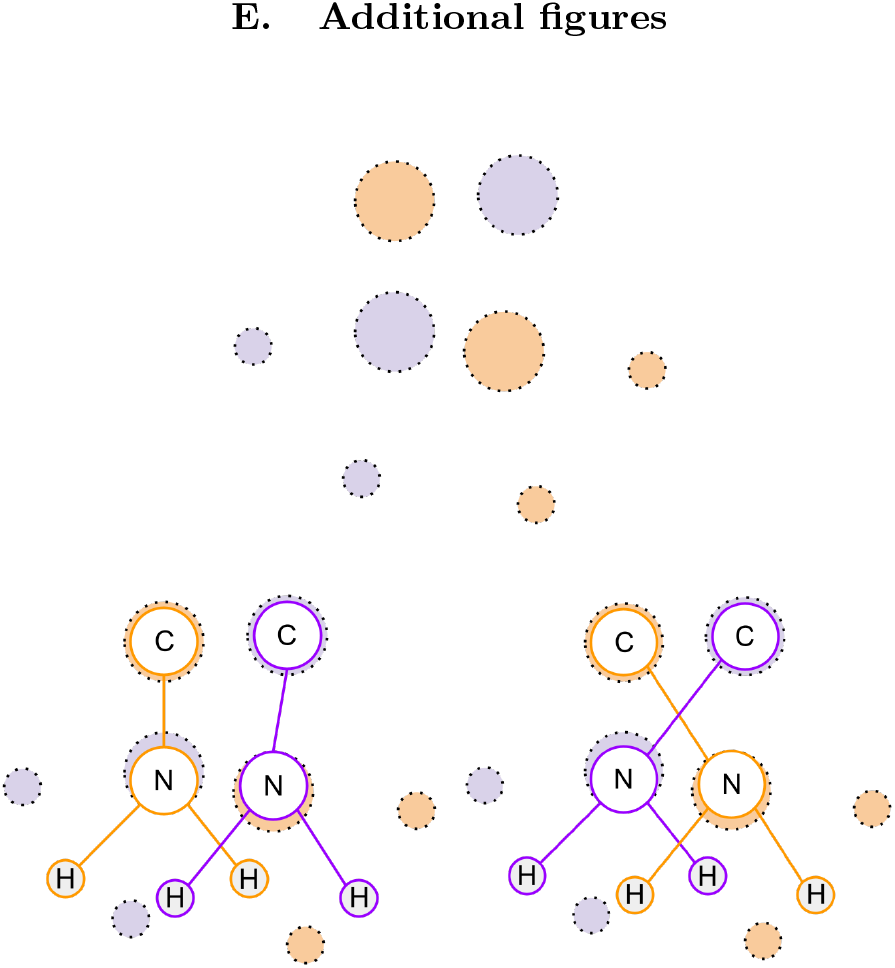
Why hydrogen geometries are ignored in connectivity optimization. Shown is an alanine side chain in a 2conformation model. Filled-in dotted-edge circles represent the density, colored according to the ground-truth conformation (top). The initial model conformations are shown bottom left, with edges colored according to the label assigned. Because the determination of hydrogen positions is substantially more reliant on geometry than for other atoms, the hydrogens are significantly displaced from their correct coordinates due to the conformation mix-up caused by the incorrect C– N bond. The bottom right shows the model after making the necessary altloc label changes to escape the conformation mix-up. While the model will relax to a good geometry after refinement, the C-N-H angles are initially far from their ideal values for both conformations, and so would carry a high penalty if considered by the connectivity optimizer.

**FIG. 14.**
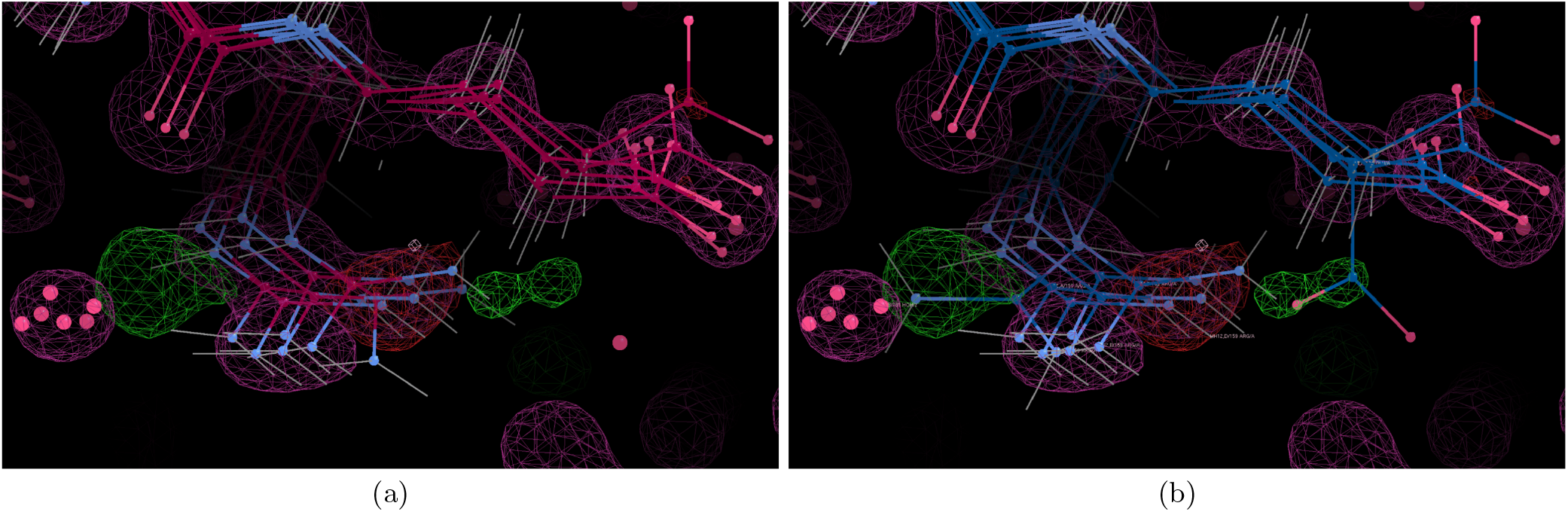
Revealing a correlated conformational difference in two adjacent side chains of the DHFR model. (a) Sidechains of Arg159 (bottom-left) and Glu134 (top-right) in the 6-conformation model after automated refinement and connectivity optimization (end of Stage 2 in Fig. 10). (b) Model after manually rotating the two side chains from right to left for one of the conformations, each replacing a partial-occupancy water. The electron density maps correspond to the model in (a) for both figures. In (a), the water altlocs left of Arg159 are not completely overlapping with density, but they are close enough as to clash with any rotamer placed to fit the adjacent positive peak (green) in the FoFc map. The positive and negative peaks at the Arg159 side chain are similar, suggestive of genuine heterogeneity rather than radiation damage.

## V. Discussion

We have introduced a procedure for resolving conformation mix-ups using “connectivity optimization” – a method to connect model altlocs into new, less-strained conformations without altering the modeled structure factors. This method was able to resolve all conformation mix-ups in the synthetic 2-conformation challenge dataset constructed by Hopkins *et al*. [20]. While work remains in tuning an automated method for practical application, including connectivity optimization in a refinement procedure produced moderate reductions in geometric strain and *R*_free_. This reduction was likely limited without an automated method of handling multiconformer errors that are unrelated to altloc connectivity. However, such errors were substantially reduced in 6-conformation models by adding low-occupancy waters to each conformation to fit density features in the bulk-solvent.

Earlier work has predicted that the gap in *R*-factors between small-molecule crystallography and macromolecular crystallography is not due to noise, but due to limitations in modeling [13]. While protein models seldom reach *R*_free_ values as low as 10–15%, the considered work showed that, even accounting for realistic sources of error in data collection, models should be able to approach *R*_free_ values close to 5%. The 6-conformation model of SARS-CoV-2 Mac1 presented here, with an *R*_work_/*R*_free_ of 4.7%/6.4%, signifies this to indeed be the case. This suggests revisiting the wealth of already deposited data will offer new structural insights [20]. Initiatives in this direction are already underway; as highlighted by the recent release of over 60,000 single-conformation PDB models re-refined as multi-conformer models with `qFit` [14].

A common criticism of ensemble models is the risk of overfitting posed by the substantial increase in parameters over single-conformation models (e.g. [10]). However, these results suggest this criticism is somewhat premature. A deposited 250-conformation model for the DHFR dataset with anisotropic *B*-factors (PDB ID: 4PTH [36]), represents orders of magnitude more parameters than the isotropic *B*, 6-conformation model presented here. As such, its *R*_work_/*R*_free_ values (12.6%/14.4%) being substantially higher than the 6-conformation model (6.3%/8.2%) demonstrates it is trapped, far from the global minimum fit to the experimental data.

The additional challenges encountered in extending the method to real data shows the scope of ensemble-specific issues identified by Hopkins *et al*. [20] expands beyond just conformation mix-ups. In the synthetic scenario, traps arise from ambiguity as to which conformation should fit which region of density; however, the identity of the atoms responsible for said density is unambiguous. As a result, the connectivity optimization approach, combined with limited coordinate change options, was sufficient to untangle the model. In contrast, solvent and protein atoms of real samples in different asymmetric units frequently make overlapping contributions to the electron density map, greatly increasing the risk of identity mix-ups. This suggests a need for ensemble-specific methods to identify potential errors in the attribution of one or more atom identities responsible for a given electron density feature.

In adding ordered waters to real models in Sec. IV B, each water was assigned one of the conformation labels used by the protein altlocs. However, the water occupancies were refined individually, independent of the protein conformations. This treatment represents a compromise between constraining water positions using non-bond interactions with protein conformations, and the need for flexibility to fit non-flat bulk-solvent features. However, this approach runs into issues of physical consistency and model interpretability. Ideally, a direct link could be drawn between the modeled conformation, and the placement of water around it. A possible solution would be to encode conformations as a heterogeneity hierarchy [44]. With this format, each water can be represented in a child conformation of a particular protein conformation; offering direct association without sacrificing the capability to model weak density features, in the same way that conventional structural models frequently contain multiconformer waters. The connectivity optimization algorithm supports solving problems with hierarchical heterogeneity, but refinement packages do not handle the format at present.

The method to correct coordinate mix-ups for atoms in two equal-occupancy altlocs was shown to be effective when applied to the synthetic scenario. The connectivity optimizer replaced altloc coordinate pairs with ones that modeled similar density but corrected connectivity issues in the main chain. While this method was not extended to handle atoms in three or more altlocs, the signature of coordinate mix-ups observed in the synthetic 2-conformation case – altlocs on the short axis of the density ellipsoid – suggests these traps are possible because multiple distinct sets of altloc coordinates can provide similar or local minima fits to a density “blob” with low geometric strain. As such, it is plausible that models with more conformations are more susceptible to coordinate mix-ups. A method to handle coordinate mixups involving more than two altlocs might resemble the core routine of `qFit`, which processes many candidate combinations of conformations for fitting local regions of density [15].

The alternative hypothesis models identified in the synthetic challenge scenario appeared to only be possible due to their singular coordinate mix-ups. It is important to note that this model still represents a substantial improvement over the starting model; with the average RMSE in the coordinates of individual conformations (by comparison with `best.pdb`) reduced from 0.13 Å to 0.084 Å. Additionally, this may be an artifact of the idealized assumptions behind the synthetic 2-conformation scenario. If two local minima states in the energy landscape did indeed differ due to a coordinate mix-up, a particular molecule might pass through both conformational states at physiological temperatures. As such, if a similar scenario were to arise in practice, it would be reasonable to model both possibilities. On the whole, it is encouraging that the only alternative hypothesis models we were able to identify represent models with substantially reduced differences from the truth compared to the (poorer metric) models converged on when refinement is trapped by tangling.

The method presented here is intended to be demonstrative. The connectivity optimization program considers bond, angle, non-bond and clash terms in its cost function, but other measures employed by modern refinement software – such as planarity and chirality – have not been implemented here. Little optimization of weights was done; only modifying the weights applied by `phenix` to uniformly increase the weight placed on angles. There is no attempt at handling genuine outliers. Ideal values for geometries are taken as single values, but standard practice is to use values that are conformation-dependent, such as with the conformation-dependent library [26]. The robustness of the method would also be improved with a treatment for swapping altlocs between atoms with differing PDB format names; particularly for symmetric groups such as the guanidino group [R-HNC(NH_2_)_2_] of arginine. More sophisticated strategies are also needed for efficiently exploring the search space of connectivity optimization in models with more conformations.

## VI. Conclusions

Ensemble models can become trapped connecting electron density that should belong to separate conformations, straining geometry. This tangling pheomenon is a connectivity issue between the altlocs of the ensemble across all conformations. It frequently manifests as locally strained geometries in distant parts of the structure, and thus is rarely apparent from inspection. We have demonstrated that these traps may be relieved through connectivity optimization by formulating the energy-minimizing assignment of altloc labels as an integer linear programming (ILP) problem. Extending the ILP to allow for multiple altloc coordinate sets for individual atoms was shown to enhance results for synthetic data.

This work builds on a thesis that the central outstanding challenge to ensemble refinement is navigating a topological landscape of many plausible fits to the average electron density. While this sometimes arises in single-conformation modeling, it is the unifying theme of conformation mix-ups, coordinate mix-ups, and identity mix-ups. Developing new methods for escaping these traps has the potential to further raise the ceiling for the accuracy achievable in macromolecular modeling. Both in representing the average structure (as measured by *R*_free_), and individual, more physiologically relevant structural states.

## Acknowledgements

We thank Dr. Alaric Sanders for his generous and thoughtful discussion on the nature of the tangling problem in the nascent stage of the project. We additionally extend our thanks to Dr. Chris Szeto, Prof. Harry Quiney, Prof. James Fraser, and Dr. Stephanie Wankowicz for their feedback on the interpretation of the 6-conformation experimental models. This research was supported by the Australian government through the Australian Research Council’s Future Fellowship funding scheme (FT220100405). We acknowledge additional funding from the Australian Research Council (DP250100311).

## Generative AI usage statement

Generative AI was not used in the research design, code implementation, or writing pertaining to this article.

## Code availability

The connectivity-optimizer code repository is available at https://github.com/Phoelionix/Untangler. Since the algorithm is under active development, the code used for generating the submitted solution to the synthetic 2-conformation challenge [20] may be found in the `2 conformer challenge solution` branch of the repository.

## A. Holton weighted energy

In application to the synthetic 2-conformation dataset, the objective function of the connectivity optimizer was modified to follow the Holton weighted energy wE [20], with the exception of clash terms. wE is similar to the sum of statistical potentials used in refinement algorithms [Eq. (2)], but with each term weighted to place a higher penalty on statistical outliers. This modification is motivated by the argument that refinement algorithms often favor models with a small number of physically impossible geometric outliers over those with a large number of moderate geometric deviations.

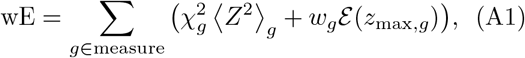

Where

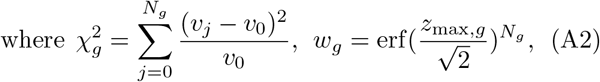

And

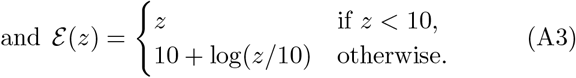

Equation A1 sums over each kind of measure *g*; for example covalent bond length, or planarity. The term *z* is the unweighted *z*-score, *z*_max,*g*_ is the *z*-score of the largest outlier for geometry measure kind *g*, and *N*_*g*_ is the number of values for the kind of measure. In Eq. A2, *v*_*j*_ is a specific instance of measure kind ⟨*g. Z*^2^⟩*g* is the average value of *Z*^2^ for measure kind *g*.

When *g* corresponds to non-bond/VDW interactions, *Z*^2^ is transformed as

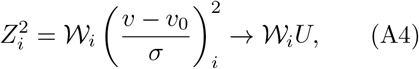

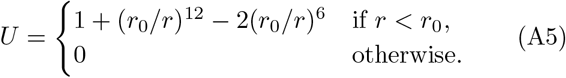

The term *U* equates to the Lennard-Jones potential, clipped when the separation *r* is above the minimum-energy separation (*r*_0_) as given in the CCP4 monomer library [22].

In Section III, the aim of the connectivity optimization stage is to minimize wE. The form of Eq. A1 is inefficient for an ILP. Instead, we use a target function with a similar form to the 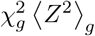 term,

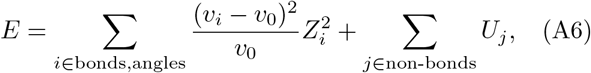

where *i* indexes all active bond, angle, and non-bond terms. Note that clashes were forbidden, so are not considered here.

### B. Untwist move detection details

The conditions where coordinate mix-ups can occur are highly constrained for 2-conformation models. In addition to the ellipsoid requirement (Fig. 6a), there should exist some altloc label assignment where the true altloc coordinates each lie roughly at the intersection of arcs that their bond and angle restraints on the atom define (Fig. 6b depicts one such arc). This does, however, assume that the altlocs of atoms within two covalent bonds are near the ground-truth coordinates, and are correctly related by their conformation labels.

Candidate untwist moves are identified by considering triplets of consecutive atoms: 1, 2, and 3. The atom altlocs are taken for the model after unrestrained refinement. The algorithm assumes that the altlocs of atoms 1 and 2 are correct. Atom 3 is considered to be in a possible coordinate mix-up if there exists a pair of coordinates for atom 3 that, consistent with an alternative hypothesis for the ground-truth altlocs,

**(I):** has approximately the same midpoint as the current altlocs,

**(II):** has greater separation than the current altlocs,

**(III):** forms angles (atoms 1–2–3) and bond lengths (atoms 2–3) that are close to their ideal values.

For condition III, changes to the altloc labels of A and B are not considered (i.e. the active bond geomections between A and B are assumed to be correct). As real geometries may deviate from ideal values, solutions are considered for angles and bond lengths halfway between their current and ideal `phenix` values.

Alternative hypothesis coordinates are generated by comparing two 25° arcs coordinates for each conformation, constrained by the angle of atoms 1–2–3 and bond length of atoms 2–3 halfway between their ideal and model values. The arc center is chosen to be at the closest point to the coordinate of the altloc of atom 3 for the given conformation. Points along the two arcs are then tested to see if they satisfy the conditions. If any points satisfy the conditions, they are separated according to whether they twist the pair clockwise or anti-clockwise, then averaged. This produces zero, one, or two pairs of alternative hypothesis coordinates for atom 3.

Where alternative hypothesis coordinate pairs are identified for the same atom from considering multiple angles, attempts are made to identify “compatible” pairs between two or more angles. Where this occurs, other alternative hypothesis coordinate pairs are removed. The remaining coordinate pairs are filtered so that no pair has an orientation within a 30° rotation of another. In application to the synthetic 2-conformation dataset (Sec. III) the number of coordinate pairs remaining for a particular atom was never more than 2.

For each possible coordinate mix-up identified, the altlocs are moved to the corresponding alternative hypothesis coordinates, and an unrestrained refinement cycle is performed to check for agreement with the X-ray data. This is repeated for atoms with more than one pair of alternative hypothesis coordinates. The coordinates were then filtered according to the changes in separation or orientation from the unrestrained refinement step. In general, significant reductions in the altloc separations relative to their original separations, and significant changes in orientation, were taken as implausible and removed from consideration. The exception to this rule was when new coordinates maintained a significant separation – more than twice the separation of the original coordinates, or more than five times the separation of the original coordinates for separations below 0.2 Å. The coordinates of the altlocs after the unrestrained refinement are passed to the connectivity optimizer as optional untwist moves. To avoid including near-duplicate moves, candidate moves were only considered when the rotation of their coordinate pair was more than 20^*°*^ relative to all previously identified candidate moves.

### C. Modified solver behavior for real data

ILP solvers can be given ‘warm starts’ wherein they are given a solution to start from. Starting with a non-optimal solution can significantly constrain the possible space for the optimal solution. In applying connectivity optimization to models with many conformations, the possible solution space of the connectivity optimization ILP is many orders of magnitude larger than when working with 2 conformations, and finding the global minimum solution may be computationally infeasible. We found that solving significantly constrained subproblems of the full ILP, then using these solutions as warm starts for later subproblems, was an effective way to significantly improve the speed at which significant model improvements could be identified.

In application to the synthetic data, we forbade activating outlier geomections except where they were closer to the ideal value than the worst geomection of the geometry. Here, we modify this behavior to require that they are lower than the worst by a certain factor. To do this, solution solving is performed over a series of 5 “rounds” where this factor decreases through 20, 4, 2, 1.01, and 0.8. Additionally, the definition of an outlier is made more stringent than in Sec. III; 2*σ* for bonds and 1.5*σ* for angles.

In each solution solving round, a subproblem is solved 5 times sequentially, considering a random subset of conformations in the ensemble. The number of conformations considered is dynamically adjusted depending on solution time. Solving ends after three minutes or when the gap between the guaranteed minimum value of *E* and the current solution for *E* is no more than 0.5%, whichever comes first. In combination with the aforementioned stepped relaxation of the solution space through consecutive rounds, this approach dramatically improved the speed of finding significant reductions in *E*.

In addition, random site and geomection switch variables of the current solution were held fixed for each sub-problem, at rates of 1 in 3000 and 1 in 5, respectively. Geomections with outliers of more than 2 standard deviations were never held fixed. The practical effectiveness of this strategy has not been tested.

### D. 6-conformation model refinement details

The initial model coordinates were deviated by an average of 0.5 Å. For each dataset, the initial model was first refined through an automated “connectivity refinement loop” consisting of one unrestrained `phenix.refine` refinement macro-cycle, connectivity optimization, and ten restrained refinement macro-cycles with a geometry weight at half the default. This was repeated until convergence in *R*_free_. Restrained refinement was performed at half the default geometry weight (`wc=0.5`). Further repeats were then performed, with conformation-aware water picking performed during the second half of restrained refinement. As `phenix.refine` sometimes adds waters without altloc labels, labels were assigned to such waters during connectivity optimization. Some waters in the Mac1 model are missing altloc labels because they were added in the final restrained refinement stage. The number of restrained macro-cycles was ten when water picking was not performed, and twelve when it was to help with convergence. Initial occupancies of new waters were set to 0.33. The minimum occupancy of waters, and their minimum separation from altlocs in other conformations, were both set to near-zero values (0.02 and 0.03 Å, respectively). The `phenix.refine` parameter `ordered_solvent.ignore_final_filtering_step` was set to `True`.

Before manual adjustments were made, hydrogens were added to the lysozyme model automatically in `coot` . The connectivity refinement loop, including water picking during restrained refinement, was run between manual adjustments typically one to three times. Water occupancies were also refined in this stage.

For the Mac1 model, *B*-factors were converted to anisotropic values, and refined with restraints until convergence. The connectivity refinement loop, ordered water addition, and manual adjustments were each performed two additional times. This was not done for the other models. Infrequently, minor deviations from the method described here were made on a case-by-case basis where justified by improvement in validation metrics [18], such as repeating restrained refinement when *R*_free_ did not converge.

Approximately thirty manual changes were made in each model, with approximately five to ten correcting issues caused by solvent fitting protein or ligand density (see below). In the case of DHFR, the total occupancy across conformations of the Mn^2+^ ions was reduced to less than half, and a chlorine ion with a total occupancy of 64% replaced a water atom. In some cases, individual residues were assigned to their own conformations, manually moved to better fit the density, then assigned back to their original conformations. Connectivity optimization would then be immediately performed in an attempt to find a connectivity that could accommodate the change. This was performed when the density supported a large change to a side chain, but doing so was intractable without producing

- strained geometry in the main chain, due to moving from one side of the main chain density to the other,
- clashes with another side chain that could be fixed with a different connectivity between the two residues, or
- clashes with waters fitting broad regions of electron density within the bulk solvent.

Main-chain connectivity optimization was performed after restrained refinement in the final loop for the lysozyme model to resolve clashes introduced by conformation-aware water picking. This was also done for the DHFR model, but manual changes and restrained refinement without water picking were subsequently performed with the water coordinates held fixed. The mainchain connectivity optimization produced two notable clashes in the DHFR model through symmetry relations between unit cells (which the connectivity optimizer does not consider) – between residues Pro21 and Gln108 in conformation E, and between residue 65Gln and a water in conformation D. These clashes may be interpreted as instances where a conformation must be adjacent to a different conformation, rather than physically implausible interactions in the model. For example, the VDW volumes of the conformations at the Pro21–Gln108 contact admits any crystal-packing arrangement where Pro21 in conformation E always neighbors Gln108 in conformation B, D, or F. As such, the aforementioned residues were held fixed alongside the waters in the final restrained refinement of DHFR.

Radiation damage was modeled in solvent-facing side chains of the DHFR structure by removing carboxyl groups from one conformation of Glu80, Glu139, and from two conformations of Glu118. Additionally, the guanidine group was removed from one conformation of Arg98. For 118Glu, this reinterprets a rotamer conformation in the original deposited model (4PSS) as a decarboxylated side chain.

### E. Additional figures

## Notes

### Competing Interest Statement

The authors have declared no competing interest.

### Summary of Updates

- Make equation 16 consistent with equation 10. - Clarify which crystal packing contacts are ignored by connectivity optimizer. - Grammar, formatting, and layout. - No longer refer to PuLP as an acronym for "Python universal Linear Programming" - this appears to be an unofficial term originating from LLM-generated text in a 2024 blog post. - Details on refinement of DHFR model in Appendix D.

